# HCN channel currents underlie distinct neurophysiology of mediodorsal thalamus subnuclei

**DOI:** 10.1101/2025.10.05.680564

**Authors:** Gregory J. Ordemann, Griffin J. Heckler, Kristen Springer, Falyne Driver, Alexander Jackson, Audrey C. Brumback

## Abstract

The mediodorsal thalamus (MD) is a hub coordinating cortical and subcortical brain regions to support executive and social/emotional functioning. MD can be subdivided into medial (M), central (C), and lateral (L) based on synaptic coupling, molecular identity, and physiology. Recently, we identified differential intrinsic properties between thalamocortical M and L neurons projecting to the medial prefrontal cortex (mPFC). L neurons projecting to mPFC showed increased hyperpolarization activated cyclic nucleotide gated (HCN) channel activity compared with M neurons, which caused L neurons to have lower cellular resistance and shorter time windows for integration of inputs. In addition to their role in synaptic integration, HCN channels are critical for thalamic rhythm generation. In this study, we used a combination of patch clamp electrophysiology and *in situ* hybridization to investigate how differences in HCN impact intrinsic oscillatory dynamics in M, C, and L neurons. We found that HCN current differed across MD subnuclei with C > L >> M. Clustering neurons based on HCN properties was sufficient to classify subnuclei with >95% accuracy, highlighting the differences in HCN function between subnuclei. Greater HCN activity in MD neurons was associated with decreased input resistance, decreased action potential firing, and higher resonant frequency. These findings provide an ionic basis for differences in cellular resonance across MD subnuclei, with implications for thalamic rhythm generation and information processing.

## Introduction

The thalamus is well-established as a center for relaying sensory information and the generation of rhythmic activity (Guillery and Sherman 2002; Llinás and Steriade 2006; Huguenard 1998; Li et al. 2017). Recent studies have elucidated regions of the thalamus that perform computations and orchestrate network-level activity to support cognitive functions (Bennett et al. 2019; Fang et al. 2025). Of these higher order thalamic subregions, the mediodorsal thalamus (MD) has been of particular interest due to its broad connectivity with prefrontal cortex (PFC) (Krettek and Price 1977; Klein et al. 2010; de Kloet et al. 2021; Lyuboslavsky et al. 2024) and its role in executive processing (Alonso and Swadlow 2015; Mitchell 2015). MD influences a range of functions such as working memory (Alexander and Fuster 1973; Fuster and Alexander 1971; Parnaudeau et al. 2013; Bolkan et al. 2017), cognition (Ouhaz et al. 2018; Rikhye et al. 2018; Hwang et al. 2020; 2022; de Kloet et al. 2021), and social behavior (Ferguson and Gao 2018; Ramesh et al. 2025). MD can be subdivided into medial (M), central (C), and lateral (L) based on synaptic coupling (Krettek and Price 1977; Klein et al. 2010), physiology (de Kloet et al. 2021; Lyuboslavsky et al. 2024), and molecular identity (Onishi et al. 2022) specific to each subnucleus.

Recently, we identified differential intrinsic properties between thalamocortical M and L neurons projecting to the medial prefrontal cortex (mPFC). M neurons projecting to mPFC showed decreased hyperpolarization activated cyclic nucleotide gated (HCN) channel activity compared with L neurons (Lyuboslavsky et al. 2024). HCN activity, in conjunction with low threshold, transiently activated voltage gated calcium channels (Ca_v_3 VGCCs) is critical for the generation of rhythmic activity in thalamic neurons (Zobeiri et al. 2019; Llinás and Steriade 2006; Kanyshkova et al. 2012). Of four HCN channel subunits (HCN1-4) HCN4 followed by HCN2 have been identified as the most prevalent in MD (Notomi and Shigemoto 2004). However, it remains unknown whether the HCN subunit distribution varies across MD subnuclei or if there are differences in expression levels of the same subunit.

Changes in HCN channel subunit expression and the density of expressed channels can result in disruptions in oscillatory activity and lead to thalamocortical dysrhythmia, a syndrome implicated in a variety of neurological and neuropsychiatric disorders (Llinás et al. 1999). The connection between thalamic rhythm generation and HCN expression makes it critical to understand how subnuclei of a single thalamic subregion can function normally despite differences in HCN activity in wild type mice. Despite a growing body of literature describing the importance of MD for complex executive function (Wolff and Halassa 2024), our understanding of how neurons from each MD subnucleus coordinates incoming signals to elicit a desired behavioral output remains an open question. Understanding how HCN activity in subnuclei of MD affects neuron function in WT neurons provides critical information for treatment of disorders stemming from abnormal rhythmic activity.

In this study we utilized a combination of current clamp and voltage clamp electrophysiology in conjunction with *in situ* hybridization to investigate how HCN impacts the intrinsic function of M, C, and L subnuclei of MD. We measured HCN current and the relationship between MD subnucleus and the effect of HCN current magnitude on physiological properties. Further, we identify properties associated with HCN that allow for the accurate determination of MD subnucleus using physiological properties. When this method of physiological identification is used in conjunction with location information, the MD subnucleus of a given neuron can be determined with a high degree of accuracy. We found that MD subnuclei had different levels of HCN current and V1/2 measurements which in turn resulted in differences in current clamp properties like input resistance, voltage sag, and membrane time constant. Voltage clamp measurements of HCN channel properties and our RNAscope assessment of HCN subunit presence in MD suggest differential levels of HCN2 subunit expression across MD that could dictate neuronal sensitivity and functional output.

## Materials and Methods

### Animals

All experiments were conducted in accordance with procedures established by the Institutional Animal Care and Use Committee (IACUC) at the University of Texas at Austin or the University of Connecticut. Animals included in this study were C57Bl6 (JAX stock #000664) adult male and female mice 8–16 weeks in age. Adult mice were group-housed in open-top cages and fed *ad libitum*. Animals were kept in reverse lighting conditions (9am–9pm dark) and were euthanized for patch clamp physiology experiments in the mornings, around the beginning of the animals’ dark cycle. There were 47 total mice used in these experiments; 25 male and 22 female. We identified no consistent differences between male and female MD neurons across subregions, however, when split into sexes our data are underpowered to draw firm conclusions on the lack of difference in intrinsic properties based on mouse sex.

### Slice preparation

Except as noted, all reagents for patch clamp electrophysiology solutions were purchased from Sigma Aldrich. Acute slices 200 *μ*m thick (Leica VT1200) were prepared from mice 8 to 16 weeks old after being anesthetized with intraperitoneal injection of ketamine / xylazine (90 / 10 mg/kg; Acor/Dechra). Mice were transcardially perfused with 20 mL of ice-cold cutting solution containing (in mM): 205 sucrose, 25 NaHCO_3_, 2.5 KCl, 1.25 NaH_2_PO_4_, 7 MgCl_2_, 7 dextrose, 3 Na pyruvate, 1.3 sodium ascorbate, and 0.5 CaCl_2_ bubbled with 95% O_2_ / 5% CO_2_. Slices were incubated in holding solution containing (in mM): 125 NaCl, 25 NaHCO_3_, 2.5 KCl, 1.25 NaH_2_PO_4_, 25 dextrose, 2 CaCl_2_, 2 MgCl_2_, 1.3 sodium ascorbate, and 3 Na pyruvate at 37±1°C for 30 minutes. Slices were then kept for at least 30 minutes at room temperature before being used for recordings.

### Intracellular recordings

Artificial cerebrospinal fluid (aCSF) contained (in mM): 125 NaCl, 25 NaHCO_3_, 12.5 D-glucose, 3 Na-pyruvate, 2.5 KCl, 2 CaCl_2_, 1.25 NaH_2_PO_4,_ and 1 MgCl_2_. Slices were continuously perfused with aCSF in an immersion chamber (Warner Instruments) with temperature maintained at 32.5 ± 1°C (Warner Instruments TC-324C). Synaptic blockers 5 *μ*M Gabazine, 100 *μ*M NBQX, and 25 *μ*M D-AP5 were added to the aCSF unless otherwise specified.

Somatic whole-cell patch recordings were obtained from neurons in the medial (MD-M), central (MD-C), or lateral (MD-L) subnuclei of WT mice, using DODT (Zen 2.5 blue addition, Zen pro) contrast microscopy on an upright microscope (Zeiss Examiner D1). Patch electrodes (tip resistance = 4.5 – 7.0 MΩ) were filled with the following (in mM): 120 K-gluconate, 10 KCl, 10 HEPES, 10 tris-phosphocreatine, 4 MgATP, and 0.3 Na-GTP (pH adjusted to 7.37 with KOH; 315-330 mOsM). Recordings were made with Clampex 10.7 software running a Multiclamp 700B (Molecular Devices). Signals were digitized at 20 kHz and lowpass filtered at 4 kHz.

After break in and membrane potential stabilization, data were collected at holding voltages of -55mV, -65mV, and -75mV. Cells were provided steady state current to maintain the membrane potential at these potentials ± 3 mV. Except for measurements of resting membrane potential, all data reported here were taken from recordings performed at these three distinct holding voltages. Series resistance was usually 10–20 MΩ, and experiments were discontinued above 30 MΩ or if action potentials failed to overshoot 0 mV. Experiments typically lasted <60 minutes total. Liquid junction potential was estimated to be 14.3 mV using Patchers Power Tools (IGORpro, Wavemetrics). Membrane potentials are reported without correcting for liquid membrane junction potential. We typically obtained 3-5 brain slices containing MD from each animal and recorded one neuron from each slice. Sample sizes are reported as the number of neurons. The data sets were sampled from 57 WT mice.

### Electrophysiological properties

#### Current Clamp

##### RMP, τ, and R_**N**_

We estimated the resting membrane potential (RMP) as the membrane voltage recorded immediately after switching to current clamp configuration. We calculated membrane time constant (τ) using a double exponential fit to the decay of voltage after a -20 pA 50 ms stimulus that was averaged over 20 repetitions.

We estimated neuronal membrane resistance (*R*_*N*_) from the steady-state voltage change measured in response to 5000 ms current steps ranging from -40 to +15 pA in 5 pA increments. We calculated R_N_ as the slope of the linear relationship between subthreshold steady-state voltage and input current.

##### Voltage sag ratio

To estimate HCN channel activity, we calculated the voltage sag in response to 1000 ms current steps ranging from -250 to +350 pA in 25 pA increments. Voltage sag ratio was calculated by dividing the steady state voltage of the last 25 ms of the step divided by the maximum level of hyperpolarization for each sweep.

##### Action potential firing

We quantified action potential firing during five second current steps from -250 to +350 pA in 25 pA increments. We estimated the action potential threshold as the point at which the third derivative of the membrane potential exceeded 0.3. FI plots were analyzed based on the linear slope of the initial increase in action potential firing and the x-intercept of the fitted line.

##### Resonant frequency (R_f_)

R_f_ was assessed using the same Chirp protocol as described above, isolated by the application of 0.5 *μ*M TTX, 20 *μ*M Mibefradil, and 20 *μ*M Nickel to remove sodium and calcium spikes. We used MATLAB to create a code that extracted R_f_ from the resulting traces.

#### Voltage Clamp

##### Normalized current

Figures 1-3 show patch clamp recordings from MD neurons in both current clamp and voltage clamp configurations using an Axopatch 700B amplifier running pClamp 10.7 software (Molecular Devices, San Jose, CA). Upon break in, current clamp measures were made at -65 mV by delivering current stimuli from -40 to 15 pA in 5 pA intervals to measure subthreshold properties and from -250 to 350 pA in 25 pA intervals to measure a broader range of subthreshold and suprathreshold properties. After completing current clamp recordings, ion channel blockers were applied to the extracellular bath saline to isolate HCN current. These blockers included (in mM): 0.001 TTX, 0.05 Ba2+, 0.2 Cd2+, 0.1 Ni2+, 10 TEA, 1 4-AP, 1 DAP, 0.02 DNQX, 0.05 AP5, 0.002 Gabazine. Neurons were monitored in voltage clamp until the ion channel blockers had taken effect, and spiking activity was no longer observed in the test pulse traces (at least 10 minutes). The capacitance of the cell was measured using a test pulse in the pClamp software. MD neurons were held at -30 mV and stepped from -50 to -140 mV in 10 mV intervals. Traces were leak subtracted using a p/-4 method and each step was performed 3 times and averaged during analysis. After measuring HCN current, 0.04 mM ZD7288 was applied to the slice and measurements were repeated. Data were analyzed using IGOR pro 7.0 (Wavemetrics, Portland, OR). HCN current traces were filtered at 1-3 Hz. Current amplitude was measured by subtracting 50 ms averages of points from the baseline and the end of each voltage step. Maximum Ih is measured at -130 mV because repeated measurements at -140 mV resulted in the degradation of patch recordings in some neurons

**Figure 1:**
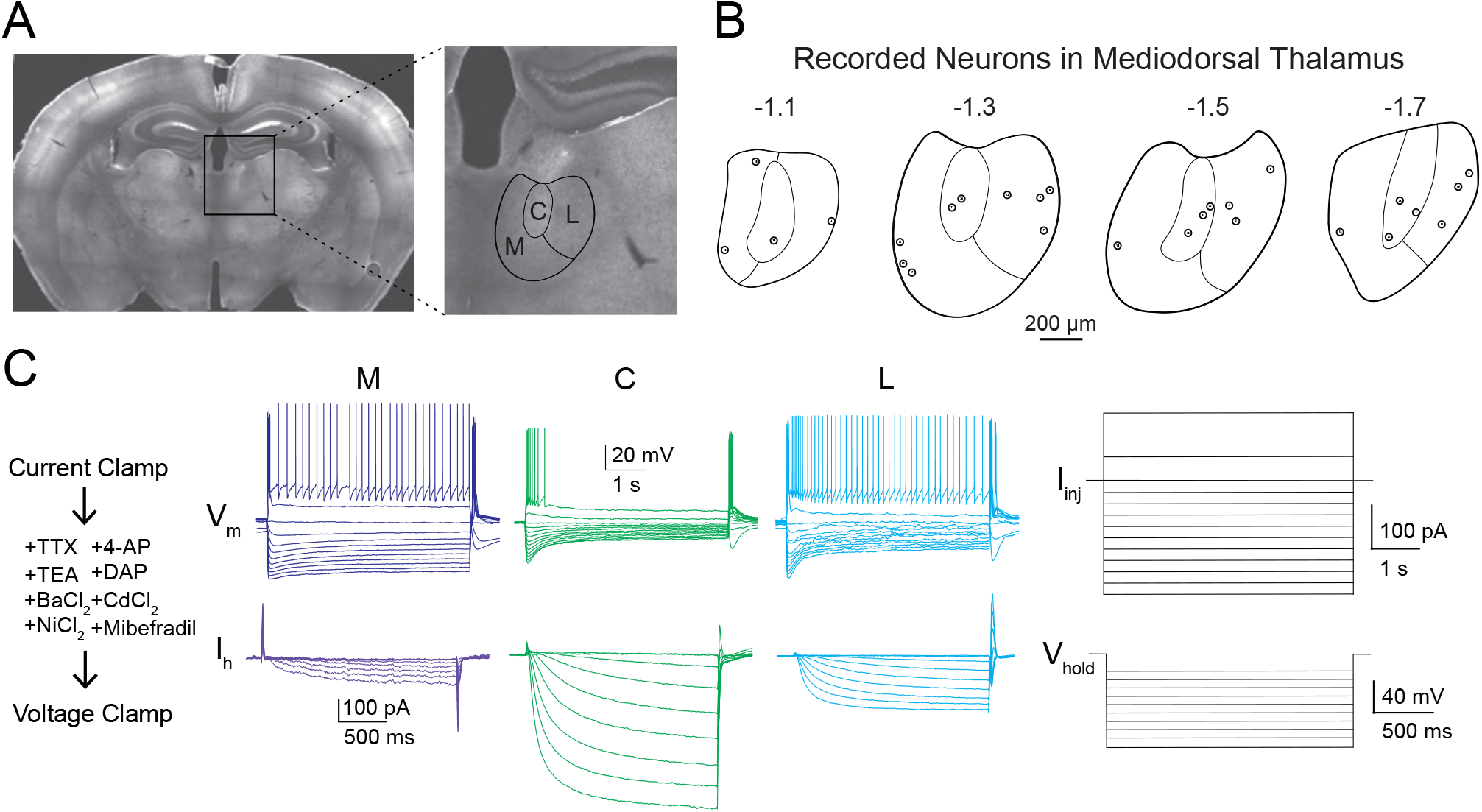
Variability in electrophysiological properties in recordings from anatomical subnuclei in the mediodorsal thalamus (MD). **A**. Left: Coronal slice of mouse brain tissue from Bregma -1.3 mm. Box shows area in zoomed image on the right. Right image shows MD and an approximation of the borders of previously delineated subnuclei (medial – M, central – C, and lateral – L). **B:** Location of MD neurons in which both current clamp and I_h_ voltage clamp experiments were performed. **C**. Representative recording showing current clamp experiment followed by the wash-on of voltage-gated channel blockers and I_h_ measurement, with example of injected stimuli for both current and voltage clamp experiments.

**Figure 2:**
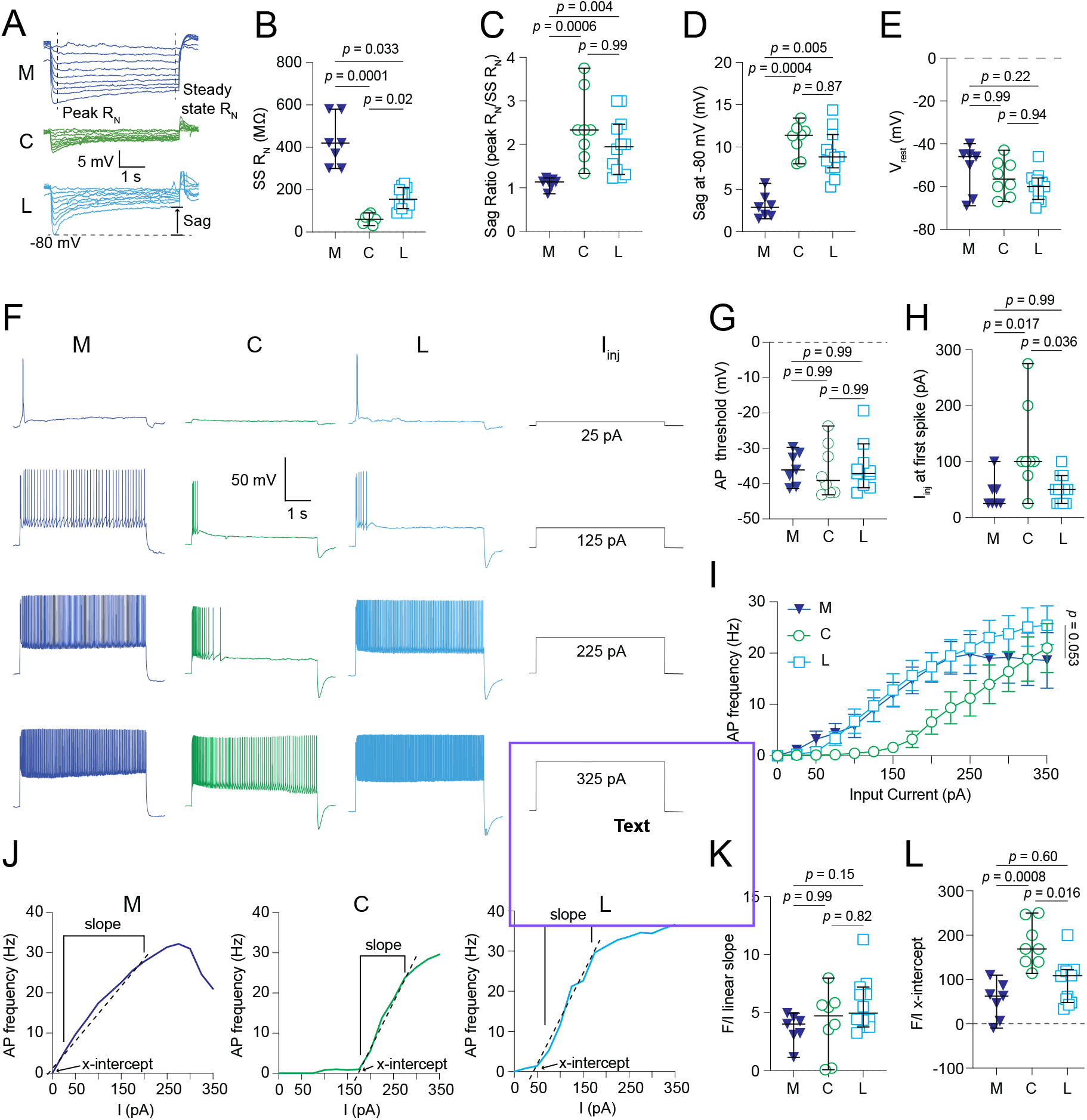
Comparison of current clamp properties across MD subnuclei. **A**. Representative traces showing voltage responses to current steps from -40 to 15 pA in 5 pA intervals. **B**. R_N_ measured as the linear fit of the steady state voltage deflection in response to the current steps from A. Kruskal-Wallis, Dunn’s Multiple Comparisons. **C**. Sag ratio measured as the quotient of peak input resistance over steady state input resistance. Kruskal-Wallis, Dunn’s Multiple Comparisons. **D**. Sag measured as the difference between steady state and peak voltage response to the hyperpolarizing current step in which the peak reached -80 mV. Kruskal-Wallis, Dunn’s Multiple Comparisons. **E**. Resting membrane potential of all cells for these experiments. Kruskal-Wallis, Dunn’s Multiple Comparisons. **F**. Representative traces showing voltage responses to current steps from +25 to +325 pA in 100 pA intervals, with accompanying input trace. **G**. Frequency of action potentials plotted against input current. **H**. Voltage threshold for neurons to fire an action potential. Kruskal-Wallis, Dunn’s Multiple Comparisons. **I**. Level of current injection in pA that elicited the first spike in each cell. Kruskal-Wallis, Dunn’s Multiple Comparisons. **J**. Frequency of action potentials plotted against input current, with slope and x-intercept measured separately for each subnucleus. **K**. Linear slope plotted from data in J. Kruskal-Wallis, Dunn’s Multiple Comparisons. **L**. X-intercept plotted from data in J. Kruskal-Wallis, Dunn’s Multiple Comparisons. **For all data**. N(neurons/animals): M=7/7, C=8/6, L=12/12

**Figure 3:**
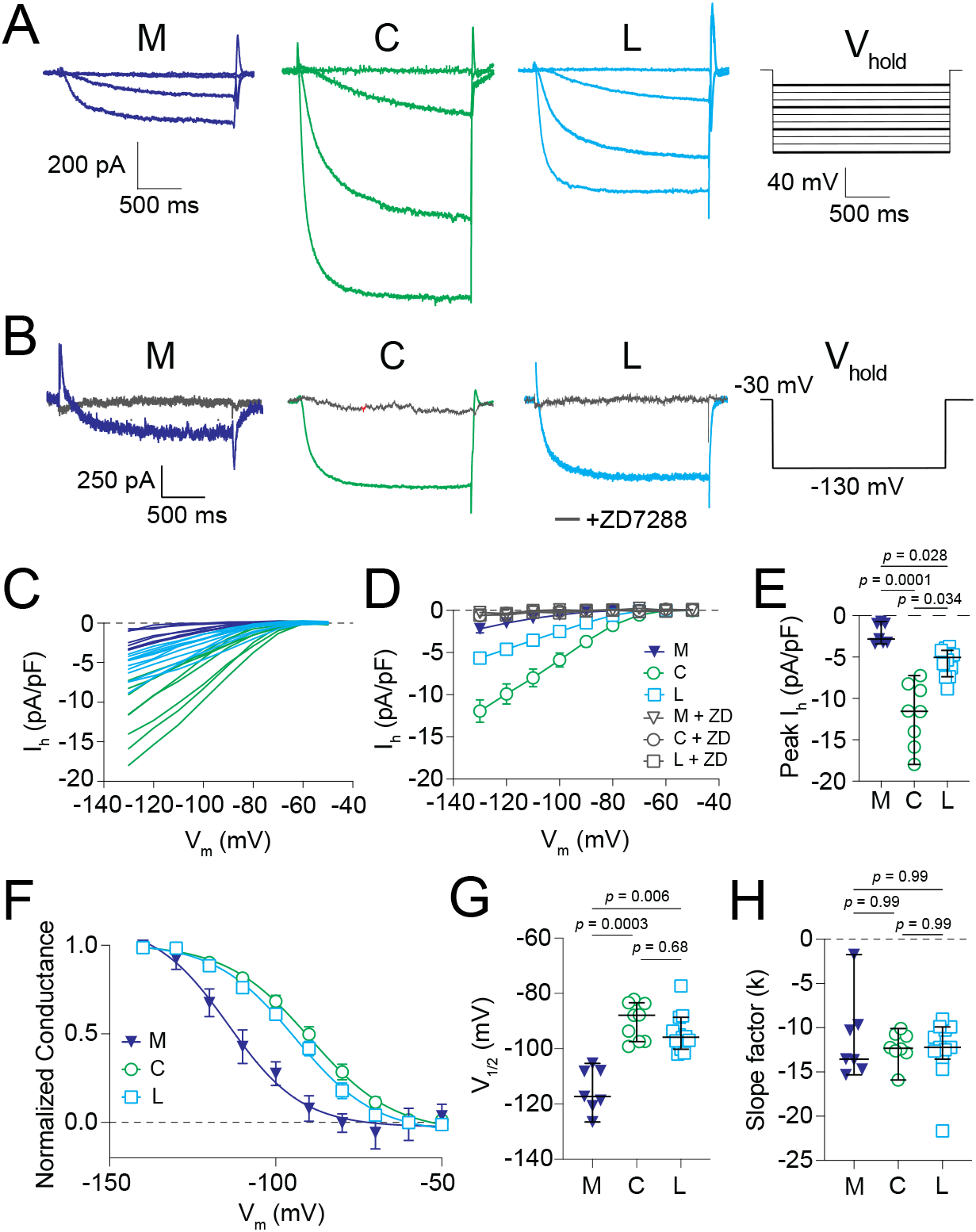
MD subnuclei show differing levels of HCN current. **A**. Representative traces from each subnucleus showing the I_h_ measured in response to steps from -30 mV to -50, -80, -100, and -140 mV. Experiment stimulus on the right with voltage steps for the present traces in thicker lines. **B**. I_h_ measurements recorded at - 30 mV holding voltage with -130 mV step before and after washon of ZD7288. Experimental stimulus on the right. **C**. I_h_ responses plotted for individual cells. **D**. Mean Ih responses for M, C, and L cells before and after ZD7288 wash-on. **E**. Peak Ih differ between MD subnuclei. Kruskal-Wallis, Dunn’s Multiple Comparisons. **F**. Normalized conductance plot showing Boltzmann fits of data representing activation curves for MD subnuclei. **G**. Voltage of half maximum activation of I_h_ for MD subnuclei. Kruskal-Wallis, Dunn’s Multiple Comparisons. **H**. Slope factor (k) for the Boltzmann fit of each neuron in F. Kruskal-Wallis, Dunn’s Multiple Comparisons. **For all data**. N(neurons/animals): M=7/7, C=8/6, L=12/12

##### Clustering

Neuron properties recorded in voltage and current clamp (Peak Ih, SS RN, peak RN, sag at -80 mV, and the xintercept of the linear fit of the F/I) were used to determine if physiological properties could be used to separate MD neurons into distinct clusters based on subnucleus. A principal component analysis (PCA) was performed and the first two components, accounting for 87.1% of variance, were used to perform the cluster analysis. The R package Nbclust (Charrad et al. 2014) was used to determine the optimal number of clusters for the data (table 1). K means clustering was performed and the clustering was validated using silhouette scores, Jaccard analysis, and adjusted rand index. Additionally, the original data set was partitioned and cross validated Jaccard and ARI tests were performed to further establish the stability of the clusters (table 1). These tests were performed in R Studio.

**Table 1.**
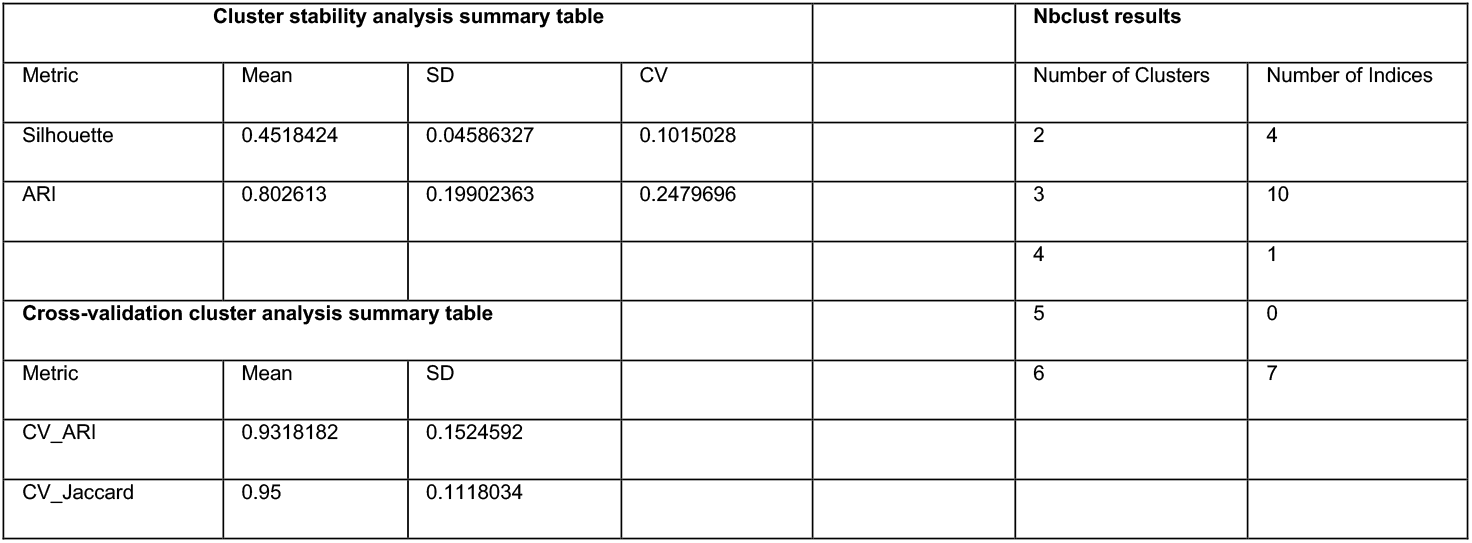
Descriptive statistics from clustering analysis.

**Table 2.**
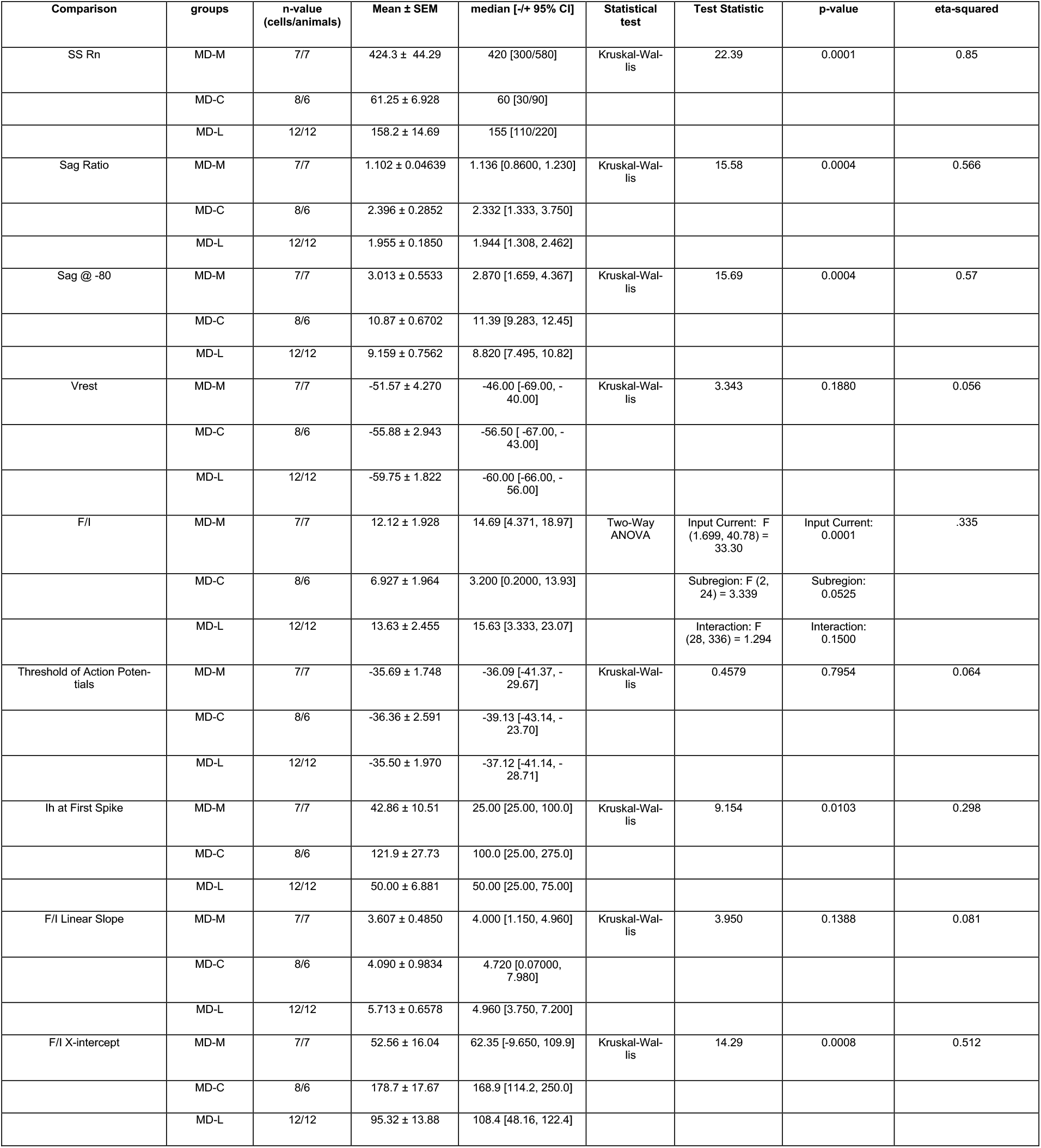
Descriptive statistics and test statistics from Figure 2.

**Table 3.** Descriptive statistics and test statistics from Figure 3.

| Comparison | groups | n-value<br>(cells/animals) | Mean ± SEM | median [-/+ 95% CI] | Statistical<br>test | Test Statistic | p-value | eta-squared |
| --- | --- | --- | --- | --- | --- | --- | --- | --- |
| Peak Ih | MD-M | 7/7 | -120.4 ± 32.70 | -147.1 [-200.4, -40.34] | Kruskal-Wal-<br>lis | 19.26 | 0.0001 | 0.719 |
| | MD-C | 8/6 | -916.7 ± 143.3 | -855.6 [-1256, -577.7] | | | | |
| | MD-L | 12/12 | -353.0 ± 42.25 | -340.5 [-445.1, -<br>261.0] | | | | |
| V1/2 | MD-M | 7/7 | -115.0 ± 2.980 | -117.3 [-122.2, -<br>107.7] | Kruskal-Wal-<br>lis | 16.46 | 0.0003 | 0.603 |
| | MD-C | 8/6 | -89.39 ± 2.333 | -87.84 [-94.91, -<br>83.88] | | | | |
| | MD-L | 12/12 | -94.35 ± 2.001 | -95.81 [-98.76, -89.95] | | | | |
| Slope factor (k) | MD-M | 7/7 | -11.28 ± 1.780 | -13.56 [-15.63, -6.925] | Kruskal-Wal-<br>lis | 0.1958 | 0.9068 | 0.075 |
| | MD-C | 8/6 | -12.22 ± 0.6351 | -12.33 [-13.72, -10.72] | | | | |
| | MD-L | 12/12 | -12.47 ± 0.9641 | -12.21 [-14.59, -10.35] | | | | |
| Peak Rn/Peak Ih | MD-M | 7/7 | 449.6 ± 56.03 | 470.0 [250.0, 660.0] | Kruskal-Wal-<br>lis | 15.86 | 0.0004 | 0.578 |
| | MD-C | 8/6 | 146.2 ± 23.90 | 139.9 [60.00, 270.0] | | | | |
| | MD-L | 12/12 | 289.5 ± 22.52 | 275.0 [259.6, 340.0] | | | | |
| SSRn/Peak Ih | MD-M | 7/7 | 414.3 ± 49.75 | 420.0 [230.0, 580.0] | Kruskal-Wal-<br>lis | 21.79 | 0.0001 | 0.825 |
| | MD-C | 8/6 | 61.25 ± 6.928 | 60.00 [30.00, 90.00] | | | | |
| | MD-L | 12/12 | 146.5 ± 14.86 | 130.0 [90.00, 200.0] | | | | |
| Sag at -80 mV/Peak Ih | MD-M | 7/7 | -3.013 ± 0.5533 | 2.870 [-5.700, -1.530] | Kruskal-Wal-<br>lis | 15.69 | 0.0004 | 0.57 |
| | MD-C | 8/6 | -10.87 ± 0.6702 | -11.39 [-13.42, -8.030] | | | | |
| | MD-L | 12/12 | -9.159 ± 0.7562 | -8.820 [-11.48, -7.540] | | | | |
| F/I x-intercept/Peak Ih | MD-M | 7/7 | 52.56 ± 16.04 | 62.35 [-9.650, 109.9] | Kruskal-Wal-<br>lis | 14.29 | 0.0008 | 0.512 |
| | MD-C | 8/6 | 178.7 ± 17.67 | 168.9 [114.2, 250.0] | | | | |
| | MD-L | 12/12 | 95.32 ± 13.88 | 108.4 [48.16, 122.4] | | | | |

**Table 4.**
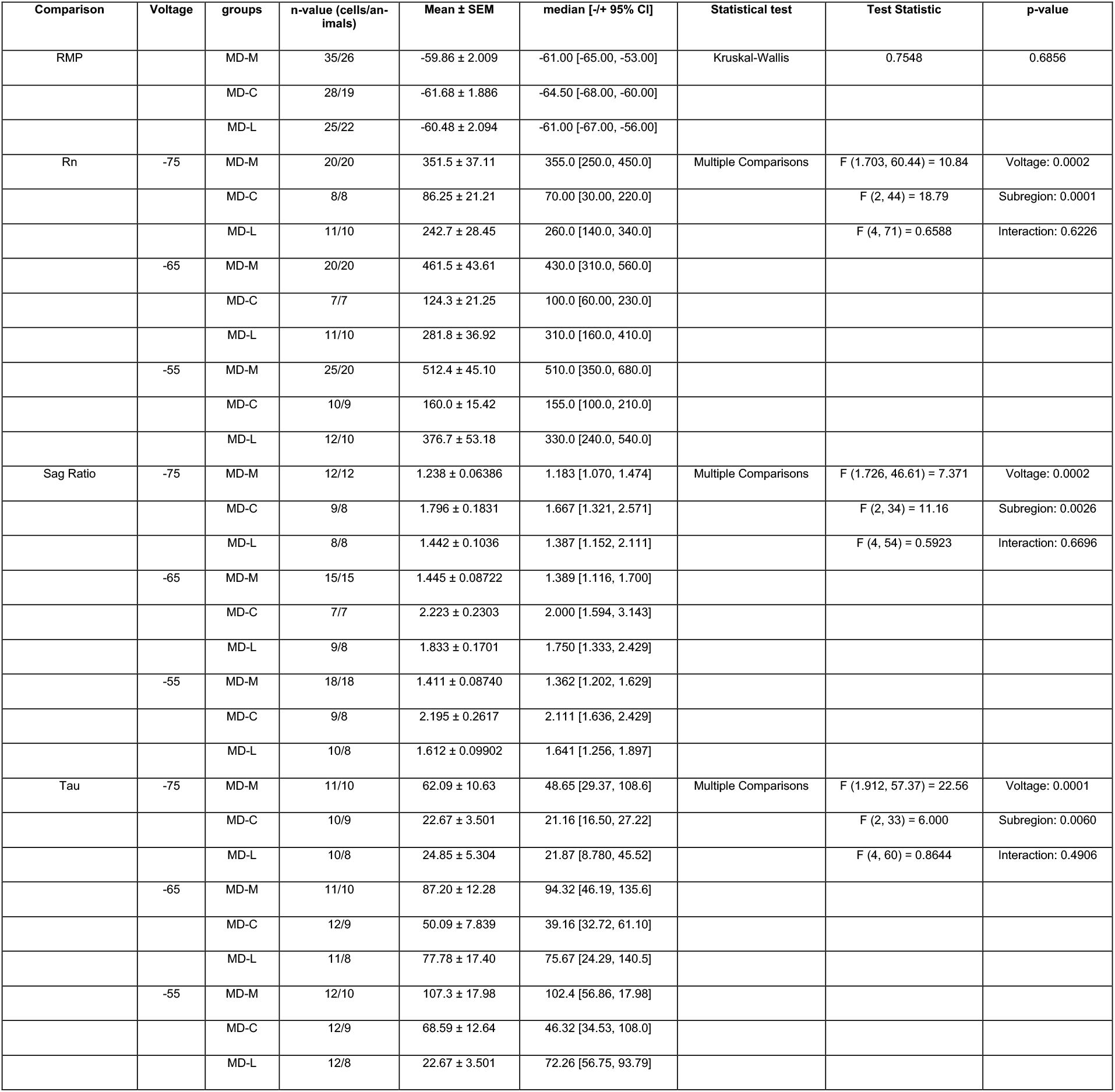
Descriptive statistics and test statistics from Figure 5.

| Comparison | Voltage | groups | n-value (cells/animals) | Mean ± SEM | median [-/+ 95% CI] | Statistical test | Test Statistic | p-value |
| --- | --- | --- | --- | --- | --- | --- | --- | --- |
| RMP | | MD-M | 35/26 | -59.86 ± 2.009 | -61.00 [-65.00, -53.00] | Kruskal-Wallis | 0.7548 | 0.6856 |
| | | MD-C | 28/19 | -61.68 ± 1.886 | -64.50 [-68.00, -60.00] | | | |
| | | MD-L | 25/22 | -60.48 ± 2.094 | -61.00 [-67.00, -56.00] | | | |
| Rn | -75 | MD-M | 20/20 | 351.5 ± 37.11 | 355.0 [250.0, 450.0] | Multiple Comparisons | F (1.703, 60.44) = 10.84 | Voltage: 0.0002 |
| | | MD-C | 8/8 | 86.25 ± 21.21 | 70.00 [30.00, 220.0] | | F (2, 44) = 18.79 | Subregion: 0.0001 |
| | | MD-L | 11/10 | 242.7 ± 28.45 | 260.0 [140.0, 340.0] | | F (4, 71) = 0.6588 | Interaction: 0.6226 |
| | -65 | MD-M | 20/20 | 461.5 ± 43.61 | 430.0 [310.0, 560.0] | | | |
| | | MD-C | 7/7 | 124.3 ± 21.25 | 100.0 [60.00, 230.0] | | | |
| | | MD-L | 11/10 | 281.8 ± 36.92 | 310.0 [160.0, 410.0] | | | |
| | -55 | MD-M | 25/20 | 512.4 ± 45.10 | 510.0 [350.0, 680.0] | | | |
| | | MD-C | 10/9 | 160.0 ± 15.42 | 155.0 [100.0, 210.0] | | | |
| | | MD-L | 12/10 | 376.7 ± 53.18 | 330.0 [240.0, 540.0] | | | |
| Sag Ratio | -75 | MD-M | 12/12 | 1.238 ± 0.06386 | 1.183 [1.070, 1.474] | Multiple Comparisons | F (1.726, 46.61) = 7.371 | Voltage: 0.0002 |
| | | MD-C | 9/8 | 1.796 ± 0.1831 | 1.667 [1.321, 2.571] | | F (2, 34) = 11.16 | Subregion: 0.0026 |
| | | MD-L | 8/8 | 1.442 ± 0.1036 | 1.387 [1.152, 2.111] | | F (4, 54) = 0.5923 | Interaction: 0.6696 |
| | -65 | MD-M | 15/15 | 1.445 ± 0.08722 | 1.389 [1.116, 1.700] | | | |
| | | MD-C | 7/7 | 2.223 ± 0.2303 | 2.000 [1.594, 3.143] | | | |
| | | MD-L | 9/8 | 1.833 ± 0.1701 | 1.750 [1.333, 2.429] | | | |
| | -55 | MD-M | 18/18 | 1.411 ± 0.08740 | 1.362 [1.202, 1.629] | | | |
| | | MD-C | 9/8 | 2.195 ± 0.2617 | 2.111 [1.636, 2.429] | | | |
| | | MD-L | 10/8 | 1.612 ± 0.09902 | 1.641 [1.256, 1.897] | | | |
| Tau | -75 | MD-M | 11/10 | 62.09 ± 10.63 | 48.65 [29.37, 108.6] | Multiple Comparisons | F (1.912, 57.37) = 22.56 | Voltage: 0.0001 |
| | | MD-C | 10/9 | 22.67 ± 3.501 | 21.16 [16.50, 27.22] | | F (2, 33) = 6.000 | Subregion: 0.0060 |
| | | MD-L | 10/8 | 24.85 ± 5.304 | 21.87 [8.780, 45.52] | | F (4, 60) = 0.8644 | Interaction: 0.4906 |
| | -65 | MD-M | 11/10 | 87.20 ± 12.28 | 94.32 [46.19, 135.6] | | | |
| | | MD-C | 12/9 | 50.09 ± 7.839 | 39.16 [32.72, 61.10] | | | |
| | | MD-L | 11/8 | 77.78 ± 17.40 | 75.67 [24.29, 140.5] | | | |
| | -55 | MD-M | 12/10 | 107.3 ± 17.98 | 102.4 [56.86, 17.98] | | | |
| | | MD-C | 12/9 | 68.59 ± 12.64 | 46.32 [34.53, 108.0] | | | |
| | | MD-L | 12/8 | 22.67 ± 3.501 | 72.26 [56.75, 93.79] | | | |

**Table 5.** Descriptive statistics and test statistics from Figure 6.

| Comparison | Voltage | groups | n-value (cells/animals) | Mean ± SEM | median [-/+ 95% CI] | Statistical test | Test Statistic | p-value |
| --- | --- | --- | --- | --- | --- | --- | --- | --- |
| Chirp Resonant Frequency | -75 | MD-M | 11/7 | 0.9465 ± 0.04849 | 0.8660 [0.8020, 1.142] | Multiple Comparisons | F (2, 60) = 5.408 | Voltage: 0.0069 |
| | | MD-C | 10/6 | 1.416 ± 0.2063 | 1.254 [0.9020, 2.082] | | F (2, 30) = 4.330 | Subregion: 0.0223 |
| | | MD-L | 12/9 | 1.246 ± 0.09219 | 1.156 [0.9500, 1.506] | | F (4, 60) = 0.9228 | Interaction: 0.4568 |
| | -65 | MD-M | 11/7 | 0.8641 ± 0.04690 | 0.8620 [0.7200, 0.9960] | | | |
| | | MD-C | 10/6 | 1.298 ± 0.1814 | 1.301 [0.7567, 1.572] | | | |
| | | MD-L | 12/9 | 0.9970 ± 0.08585 | 0.9910 [0.7675, 1.204] | | | |
| | -55 | MD-M | 11/7 | 0.8529 ± 0.04219 | 0.8433 [0.6700, 1.002] | | | |
| | | MD-C | 10/6 | 1.044 ± 0.1429 | 0.9230 [0.6320, 1.682] | | | |
| | | MD-L | 12/9 | 0.9853 ± 0.09806 | 0.9930 [0.6120, 1.172] | | | |
| ZD- Chirp Resonant Frequency | -75 | MD-M | 3/2 |  |  |  |  | MD-M vs MD-M ZD: 0.9613 |
|  |  | MD-C | 5/3 |  |  |  |  | MD-C vs MD-C ZD: 0.0061 |
|  |  | MD-L | 6/4 |  |  |  |  | MD-L vs MD-L ZD: 0.0020 |
|  | -65 | MD-M | 3/2 |  |  |  |  | MD-M vs MD-M ZD: 0.1728 |
|  |  | MD-C | 5/3 |  |  |  |  | MD-C vs MD-C ZD: 0.0300 |
|  |  | MD-L | 6/4 |  |  |  |  | MD-L vs MD-L ZD: 0.0060 |
|  | -55 | MD-M | 3/2 |  |  |  |  | MD-M vs MD-M ZD: 0.7460 |
|  |  | MD-C | 5/3 |  |  |  |  | MD-C vs MD-C ZD: 0.4341 |
|  |  | MD-L | 6/4 |  |  |  |  | MD-L vs MD-L ZD: 0.0122 |

##### Fluorescence *in situ* hybridization

Mice were deeply anesthetized with isoflurane, then decapitated. Brains were removed and immediately frozen using dry ice. Frozen brains were embedded in cryo-embedding media (OCT).14um serial sections through the medial dorsal nucleus of the thalamus were taken and directly mounted onto SuperFrost Plus microscope slides. Mounted tissue sections were then fixed using 4% paraformaldehyde (PFA) at 4°C for 15 minutes. After fixation, tissue sections were dehydrated in increasing concentrations of ethanol (50, 70, and 100%) for five minutes each at room temperature. All experiments were done using the RNAscope Multiplex Fluorescent Reagent Kit V2 (Advanced Cell Diagnostics cat# 323100) following the company protocol. Single-plex FISH was then carried out using probes against *Hcn1* (cat# 423651), *Hcn2* (cat# 427001), *Hcn3* (cat# 551521), *Hcn4* (cat# 421271), and *Gbx2* (cat# 314351). TSA Vivid 650 fluorophores were applied at a concentration of 1:1500 to all sections. All sections were then counterstained with DAPI and coverslips were placed after adding Prolong Gold Antifade Mountant (Invitrogen cat# P36930). All images were taken on a Leica Thunder imager microscope using the 20x objective. Images were then processed in ImageJ. All anatomic map overlays were acquired from the Paxinos and Franklin mouse brain atlas and adjusted to fit experiment ISH images in Adobe Illustrator.

Single plex images of the same anatomical region (approximately -1.58mm Bregma) were overlayed in ImageJ by first using the DAPI signal in each image to outline approximately the same region containing the dentate gyrus of the hippocampus and the thalamus. Once proper alignment was ensured, a five-channel composite containing signal for all *Hcn* channels and *Gbx2* was created. Intensities were measured by first drawing a line starting just outside of the centrolateral thalamic nucleus (CL) on the left side of the section and ending in about the same location on the right side to ensure complete coverage of the mediodorsal nucleus of the thalamus (MD). This line was also positioned just ventral to the paraventricular thalamic nucleus (PV) to ensure passage through each subregion of the MD. Line thickness was set to 600 pixels then intensity was measured in each channel using the ‘Plot Profile’ function. Intensity values for each channel were then moved to Igor Pro where values for each mouse were averaged, smoothed using a Savitsky-Golay filter, and normalized to the peak intensity for each channel to create the normalized intensity plot. Raw intensity values were smoothed for each animal before being plotted in Prism as mean +/SEM. Regions representing the CL on the left and right of the slice IMD were highlighted based on the *Hcn1* signal, and the area within these regions was considered the MD.

#### Statistics

We used G*Power to calculate sample sizes based on preliminary data. For patch clamp electrophysiology experiments, we estimated that between MD-M, MD-C, and MD-L, to detect a difference in membrane time constant of 25% with a standard deviation of 20 ms, given α of 0.05 and power of 0.8, we required at least 16 cells per group. N-values are reported as “neurons (number of mice)”.

We used the non-parametric Mann-Whitney test to compare two groups or Wilcoxon sign-rank test to compare paired data. To test significance of differences between groups in action potential firing as a function of input current, we used two-way ANOVA with Sidak’s test to correct for multiple comparisons. Data compared with a two-way ANOVA are reported as mean ± standard error. Data compared with non-parametric tests are presented with all data points and a single line representing the median of the data set and error bars showing 95% confidence intervals. Values for groups analyzed with nonparametric tests are reported as the median and 95% confidence intervals in the form: median, 95% CI [lower, upper]. We defined ? ≤ 0.05 with * p < 0.05; ** p < 0.01; *** p < 0.001; **** p < 0.0001. All p-values for Kruskal-Wallis, Dunn’s Multiple Comparisons are included in the figures and text. Only significant p-values for Mixed Effects, Tukey’s Multiple Comparisons test are included in the figures and text. All p-values can be found in accompanying data table. Effect size is expressed as *η*^2^, a measure of the variance in a data set arising from differences between groups. Effect size was calculated only when p<0.05. *η*^2^ effect sizes are defined as small – 0.01, medium – 0.06, and large – 0.14 (20).

Most electrophysiological analyses were performed using the Matlab software package Elecfex (Ma et al. 2024). Tau was calculating using the double exponential fitting function in Igor Pro. R_f_ was analyzed using custom Matlab code available via Github. All statistical analyses except for power analysis were performed using GraphPad Prism version 8.0.0 (GraphPad Software, San Diego, California USA, www.graphpad.com). Graphs of group data were made using GraphPad Prism and transferred to Adobe Illustrator version 24.3 for final presentation.

## Results

To investigate the differences in intrinsic properties across MD subnuclei, we made whole cell current and voltage clamp recordings of neurons in the MD region of the thalamus in acute *ex vivo* coronal mouse brain slices. Recordings were made in the MD region of the thalamus in either the M, C, or L subnuclei based on location (**Fig. 1A)**. Current and voltage clamp recording data were made between bregma -1.1 and -1.7 mm. In **Fig. 1B** pipette location of each recording is represented by black circles. Cells were patched in current clamp, then transitioned to voltage clamp after washing on synaptic blockers and voltage gated ion channels to isolate HCN channel activity (see methods; Hewitt et al. 2021). **Figure 1C** shows representative data from current clamp (top) and voltage clamp HCN channel recordings (bottom) for M, C, and L neurons.

### Subthreshold and suprathreshold voltage properties vary significantly between MD subnuclei

We measured subthreshold voltage properties by delivering a series of 5-second current steps ranging from -40 to +15 pA in 5 pA intervals and recording the responses in M, C, and L neurons (**Fig. 2A**). Previous investigations into MD intrinsic properties used 1-second current injection (Lyuboslavsky et al. 2024; Ordemann et al., n.d.). In this study we extended the injection duration to 5-seconds to capture the full decay of voltage after initial current injection to insure the capture of an accurate steady state before the offset of the current stimulus. We found significant differences between all subnuclei in steady state input resistance (**Fig. 2B**, Kruskal-Wallis: p = 0.0001), and significant differences in M versus C and M versus L neurons for voltage sag ratio (**Fig. 2C**, Kruskal-Wallis: p = 0.0004) and voltage sag at -80 mV (**Fig. 2D**, Kruskal-Wallis: p = 0.0004). Resting membrane potential did not differ significantly between subnuclei (**Fig 2E**, Kruskal-Wallis: p = 0.1880). These results support previous studies showing differences in the intrinsic properties between M and L subnuclei of MD (Lyuboslavsky et al. 2024; de Kloet et al. 2021), and further expand these results to show distinct subthreshold intrinsic characteristics between all subnuclei of MD. We further investigated the effects our observed differences in subthreshold properties have on suprathreshold properties.

To measure differences in suprathreshold properties, we delivered a series of 5-second current steps ranging from +25 to +350 pA in 25 pA steps in the same set of neurons (**Fig 2F**). The threshold for action potentials, measured from the first action potential to occur at any current level, did not differ between subnuclei (**Fig. 2G**, Kruskal-Wallis: p = 0.7954). However, the amount of current needed to elicit the first spike in these cells was significantly higher in C cells compared M and L cells as expected given the observed differences in SS R_N_. (**Fig. 2H**, Kruskal-Wallis: p = 0.0103). The frequency of action potentials across current stimulus was not significantly different between subnuclei (**Fig. 2I**, Two-Way ANOVA: Subnucleus - p = 0.0525). We further analyzed the. FI plot by measuring the slope and x-axis intercept point of the linear portion of each neuron to identify differences in rate of increasing in firing as related to current and the current at which action potential firing begins (**Fig. 2J**). The linear slope did not differ between subnuclei (**Fig. 2K**, Kruskal-Wallis: p = 0.1388). X-intercept was different across subnuclei, C was had a greater x-intercept on average than M or L indicating a right shift in FI curve (**Fig. 2L**, Kruskal-Wallis: p = 0.0008). These results indicate that subthreshold changes in neuronal properties across MD subnuclei produce an increased current, but not voltage, threshold in C neurons compared with M and L.

### HCN channel activity influences current responses differentially between MD subnuclei

Based on previous findings we hypothesized that differences in HCN channel activity underlies the subthreshold differences in intrinsic properties between M, C, and L MD neurons (Lyuboslavsky et al. 2024). After current clamp recordings were completed, synaptic and voltage gated channel blockers were applied to MD neurons to isolate HCN current. HCN current was measured by recording current responses to steps from -50 to -130 mV in 10 mV intervals from a holding voltage of -30 mV using a p/4 leak subtraction method to separate membrane current from channel current (**Fig. 3A)**. Experiment stimulus on the right in Fig. 3A shows voltage steps for the representative traces in bold lines. We confirmed that the recorded current was due to HCN channel activity by washing-on the HCN channel blocker ZD7288 at the end of voltage clamp recordings. If the current being recorded was from HCN channels then the subtracted current would be abolished, which is what we observed when comparing steps from -30 t0 -130 mV before and after ZD7288 wash-on (**Fig. 3B**). HCN currents were adjusted to the capacitance of the cell (measured using Clampex software). HCN currents across MD subnuclei increase with hyperpolarization (as expected with HCN current) and vary widely (**Fig. 3C-D**). Group data showing a lack of current after ZD7288 wash-on confirms the identity as HCN channels (**Fig. 3D**). When peak I_h_ (measured from the step to -130 mV) was compared between M, C, and L MD subnuclei we found a significant difference between groups and each subnucleus was significantly different from the other MD subnuclei (**Fig. 3E**, Kruskal-Wallis: p = 0.001). We fit a Boltzmann curve to the current response of each neuron at each recorded voltage to measure the voltage dependence of HCN channel activation (**Fig. 3F**). V_1/2_ was significantly different between subnuclei, with C and L having a more depolarized V_1/2_ of activation compared with M (**Fig. 3G**, Kruskal-Wallis, p = 0.0003). The rate of change in Boltzmann slope due to voltage, slope factor (k), was not different across MD subnuclei (**Fig. 3H**, Kruskal-Wallis, p = 0.9068). Our voltage clamp recordings reveal significant differences in the HCN channel activity across MD subnuclei. Interestingly, we also found a difference in V_1/2_ of >20 mV between C and M as well as between L and M, suggesting potential differences in subunit composition or expression levels.

To identify the relationship between current clamp measurements and the observed differences in HCN current across MD subnuclei we chose key parameters representing current clamp measurements that reflect HCN activity and plotted them against peak I_h_ adjusted to cell capacitance (**Supp. Fig. 1A-D**). We noticed that the current clamp measures in Figures 3I-L separated consistently into groups reflective of anatomical subregion when plotted against peak I_h_. We sought to identify electrophysiological measures from current and voltage clamp experiments that could be used to categorize neurons into the appropriate subnucleus.

Using variables from the cluster analysis we sought a method of identifying the subnucleus a neuron belongs to using values easily measured from current clamp electrophysiological recordings. By dividing SS RN and sag measured at -80 mV we were able to reliably separate M, C, and L cells into statistically distinct groups (**Fig. 1E, F;** Kruskal-Wallis: p = 0.0001). This method, when coupled with anatomical localization of the recording, provides a cross-validated method of identifying the subnucleus of MD neurons.

### PCA and cluster analysis of MD electrophysiological properties reveal 3 clusters that reflect the three MD subnuclei

The salient differences between MD subnuclei we observed in electrophysiological properties associated with HCN activity in both voltage and current clamp recordings led us to investigate the potential for identifying the subnucleus from which an MD neuron recording originates based on physiological properties. We performed a principal component analysis on 5 variables associated with HCN activity: peak Ih, SS RN, Peak RN, sag at -80 mV, and the x-intercept of the linear fit of the F/I. The overall contribution of each variable was comparable across the first two principal components (**Fig. 4A**). In the PCA the first two principal components accounted for 87.1% of the variance within the 5 variables provided (**Fig. 4B**). Neurons in which both current clamp and voltage clamp recordings were performed were included in a K means cluster analysis. Using the R package NBclust (Charrad et al. 2014), three clusters was the most commonly identified ideal number of clusters for this data set (Table 1). Cluster groups from the K means clustering are displayed in **Figure 4C**. One cell identified as being in the L region upon recording was clustered into the C group and is shown as a blue square within the green region representing C neurons. We validated the cluster analysis using several metrics that measure how well the data cluster within the constraints of the analysis. Silhouette values range from -1 to 1 (1 being directly adjacent to the centroid) and represent the distance of a neuron from the centroid of the assigned group. The average silhouette score was 0.43 (**Fig. 4D**). Table 1 shows additional methods of validation. The adjusted Rand index (ARI) was also used which identifies whether clustering was due to random chance. Both metrics show stable clusters that are not due to random chance. Finally, we performed cross-validated Jaccard and ARI analyses in which the initial data set was split into folds and tested as training sets and test sets of data. This method also showed high cluster stability for the cluster analysis of these data.

**Figure 4:**
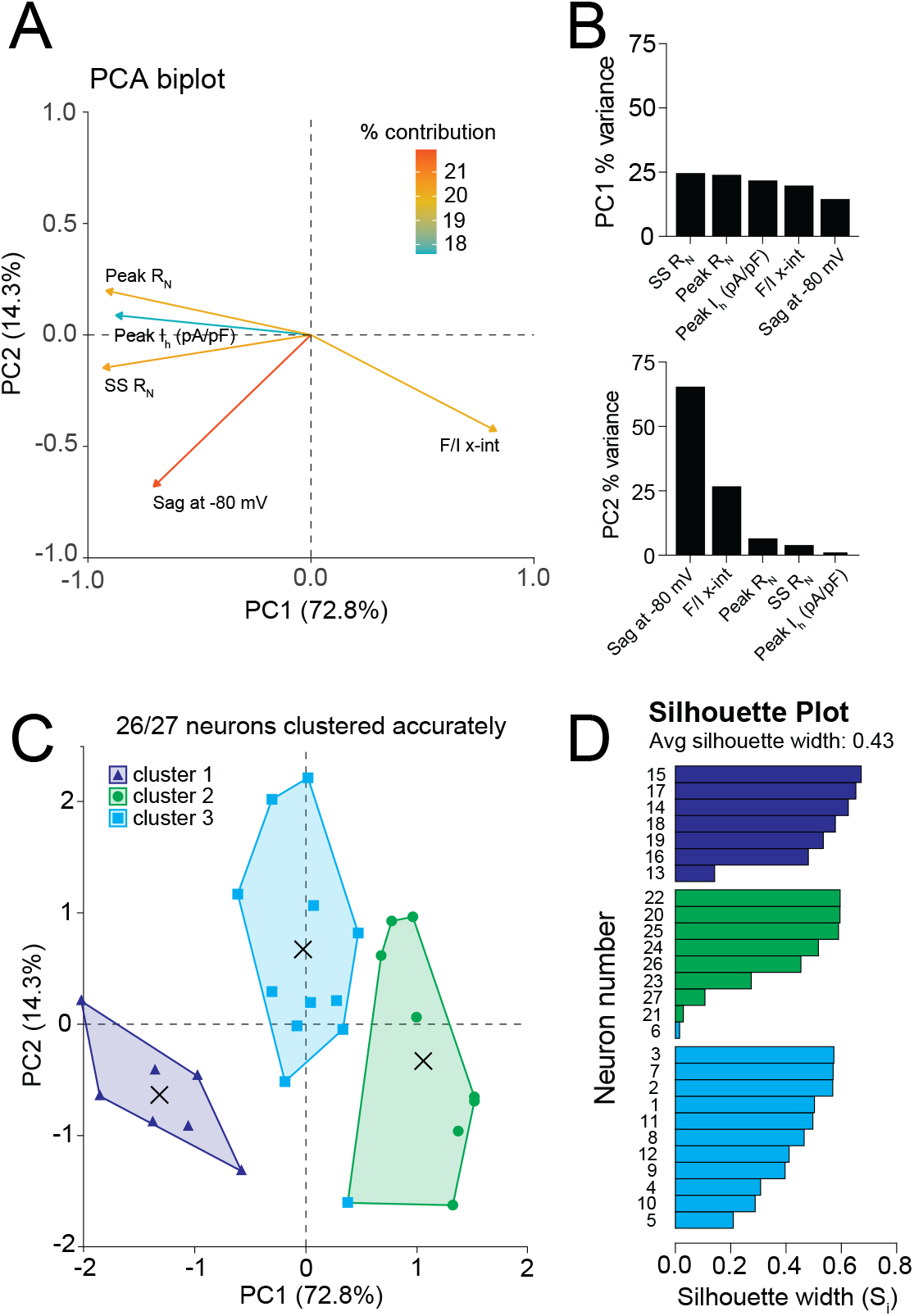
MD neurons can be identified and separated into clusters based on HCN associated properties. **A**. PCA biplot showing the contribution of each variable to the total variance accounted for by the first two principal components. **B**. Percent contribution of each variable to the first (top) and second (bottom) principal components of the PCA. **C**. K means clustering grouping results. One neuron did not cluster according to the anatomical location identified at the time of recording and is shown as a blue square within the green area of the cluster 2 (C) grouping. **D**. Silhouette scores representing how close each neuron within a cluster falls in relation to the centroid of the grouping. **For all data**. N(neurons/animals): M=7/7, C=8/6, L=12/12

#### Subthreshold properties of MD neurons differ based on membrane potential and subnucleus

To measure how subthreshold properties in MD cells vary across membrane potential, we recorded responses to 5-second current injections of -250 to 0 pA in 25pA steps at holding voltages of -75 mV, -65 mV, and -55 mV (**Fig. 5A**). Resting membrane potential for these cells did not differ significantly between subnuclei (**Fig. 5B**, Kruskal-Wallis: p = 0.8035). Input resistance varied significantly between subnuclei across holding voltages (**Fig. 5C** left, Mixed effects model: Voltage - p = 0.0002, Subnucleus - p < 0.0001, Interaction - p = 0.6226). Tukey’s Multiple Comparisons revealed differences between M and C cells at all voltages and a significant difference in RN between M and L when cells were held at -65 mV (**Fig. 5C** right). Sag ratio varied significantly between subnuclei across holding voltages (**Fig. 5D** left, Mixed Effects Model: Voltage - p = 0.0002, Subnucleus - p = 0.0026, Interaction - p = 0.6696). Tukey’s Multiple Comparisons revealed significant differences between M and C neurons at all membrane voltage levels (**Fig. 5D** right). Membrane tau was measured by delivering a 10 ms, 20 pA stimulus and averaging 20 responses at -55, -65 and -75 mV. The decay of signal after the offset of the current stimulus was fitted with a double exponential to determine the membrane tau (**Fig 5E**). Membrane tau differed across MD subnuclei with significant effects of both voltage and subnucleus. (**Fig. 5F** left, Mixed Effects Model: Voltage - p = 0.0060, Subnucleus - p = 0.0001, Interaction - p = 0.4906). Tukey’s Multiple Comparisons revealed differences in membrane tau between M and C cells at -65 and -75 mV as well as between M and L cells at -75 mV. These findings suggest that as MD neurons hyperpolarize subthreshold differences between subnuclei tend to become more salient, as is expected in the presence of differences in HCN channel activity. We next investigated how differences in HCN channel expression affect membrane oscillations in across MD subnuclei.

**Figure 5:**
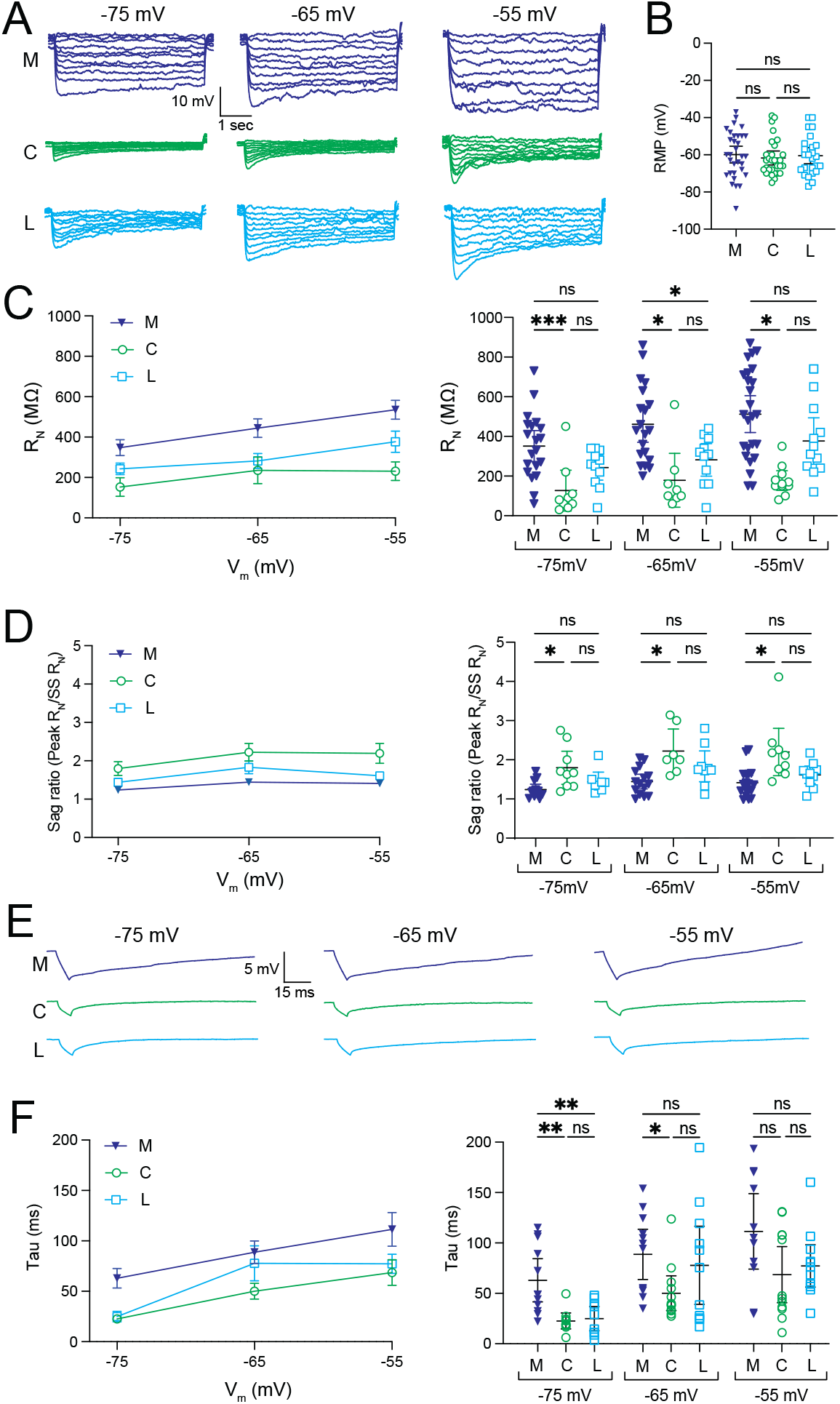
Subthreshold properties across MD subnuclei. **A**. Representative traces showing voltage responses to current step from -250 to 0 pA in 25 pA steps at holding voltages of -75mV, -65mV, and -55mV. **B**. Resting membrane potentials of MD neurons between subnuclei. Kruskal-Wallis, Dunn’s Multiple Comparisons. N(neurons/animals): M=35/26, C=28/19, L=25/22. **C**. Input resistance between MD subnuclei neurons at -75mV, -65mV, and -55mV. Data are presented with individual values in left graph and mean with standard error on right graph. Mixed Effects Model, Tukey’s Multiple Comparisons. N(neurons/animals): M=25/20, C=10/9, L=12/10. **D**. Sag ratio between MD subnuclei neurons at -75mV, -65mV, and -55mV. Data are presented with individual values in left graph and mean with standard error on right graph. Mixed Effects Model, Tukey’s Multiple Comparisons. N(neurons/animals): M=18/18, C=9/8, L=10/8. **D**. Representative traces of voltage responses to a -10 pA stimulus in MD neurons at -75 mV, -65 mV, and -55 mV. **E**. Membrane tau between MD subnuclei neurons at -55mV, -65mV, and -75mV. Data are presented with individual values in left graph and mean with standard error on right graph. Mixed Effects Model, Tukey’s Multiple Comparisons. N(neurons/animals): M=12/10, C=12/9, L=12/8

#### Subthreshold Chirp properties vary across MD subnuclei

The generation and maintenance of oscillatory activity is a critical role of the thalamus. Particularly in MD rhythmic activity in the theta to beta range has been identified as critical in working memory tasks (Parnaudeau et al. 2013). HCN channels contribute to a neurons ability to maintain rhythmic activity, acting as a band pass filter in conjunction with the electrical properties of the membrane to amplify specific frequencies (Mishra and Narayanan 2025). To determine the role of HCN channels in establishing a R_f_ in MD neurons, we washed on synaptic blockers (0.5 *μ*M TTX, 20 *μ*M Mibefradil, and 200 *μ*M NiCl_2_) to block sodium and calcium spikes. Previous studies have indicated that HCN 2 and 4 are the most prevalent subunits in MD (Notomi and Shigemoto 2004). Chirp stimuli have largely been used to assess R_f_ in cells expressing primarily HCN1 subunits which are the most voltage sensitive and have the most depolarized activation properties (Ishii et al. 2001; Santoro et al. 2000). Expecting the oscillatory properties of MD neurons to be slower than cells assayed using a linear increase in chirp frequency, we injected a chirp current stimulus of 0.5 Hz – 20 Hz that increased on a logarithmic scale tailored to expand cycles at lower frequencies. These experiments were performed at holding voltages of -75, - 65, and -55 mV. **Figure 6A-C** show the cellular responses to this stimulus by M, C, and L neurons, respectively, before and after the wash-on of ZD7288. **Figure 6D-F** demonstrates the impedance plots for all recordings from these cells plotted against the frequency of the chirp stimulus. M cells showed no deviation in resonance frequency from traces in which ZD7288 was applied, indicating a lack of resonance imparted by HCN channels (**Fig. 6G**; Two-Way ANOVA: Voltage – p = 0.68, +/-ZD7288 – p = 0.061, Interaction – p = 0.58). Both C and L neurons showed resonant properties associated with HCN channel activity since their R_f_ values diverged from ZD7288 wash-on data with C being significantly different from ZD7288 data at -65 and -75 mV (**Fig. 6H**; Mixed Effects Model: Voltage – p = 0.25, +/-ZD7288 – p = 0.011, Interaction – p = 0.11), and L differing significantly from ZD7288 wash-on data at -75 mV (**Fig. 6I**; Mixed Effects Model: Voltage – p =0.019, +/-ZD7288 – p =0.0009, Interaction – 0.012). When compared across subnuclei, it was found that R_f_ differed significantly across MD subnuclei (**Fig. 6J**; Two-Way ANOVA: Voltage – p = 0.0001, Subnucleus – p = 0.006, Interaction – p = 0.013). Sidak’s Multiple Comparison analysis revealed that C and M were different at -65 mV and C and L were different from M at -75 mV (**Fig. 6K**).

**Figure 6:**
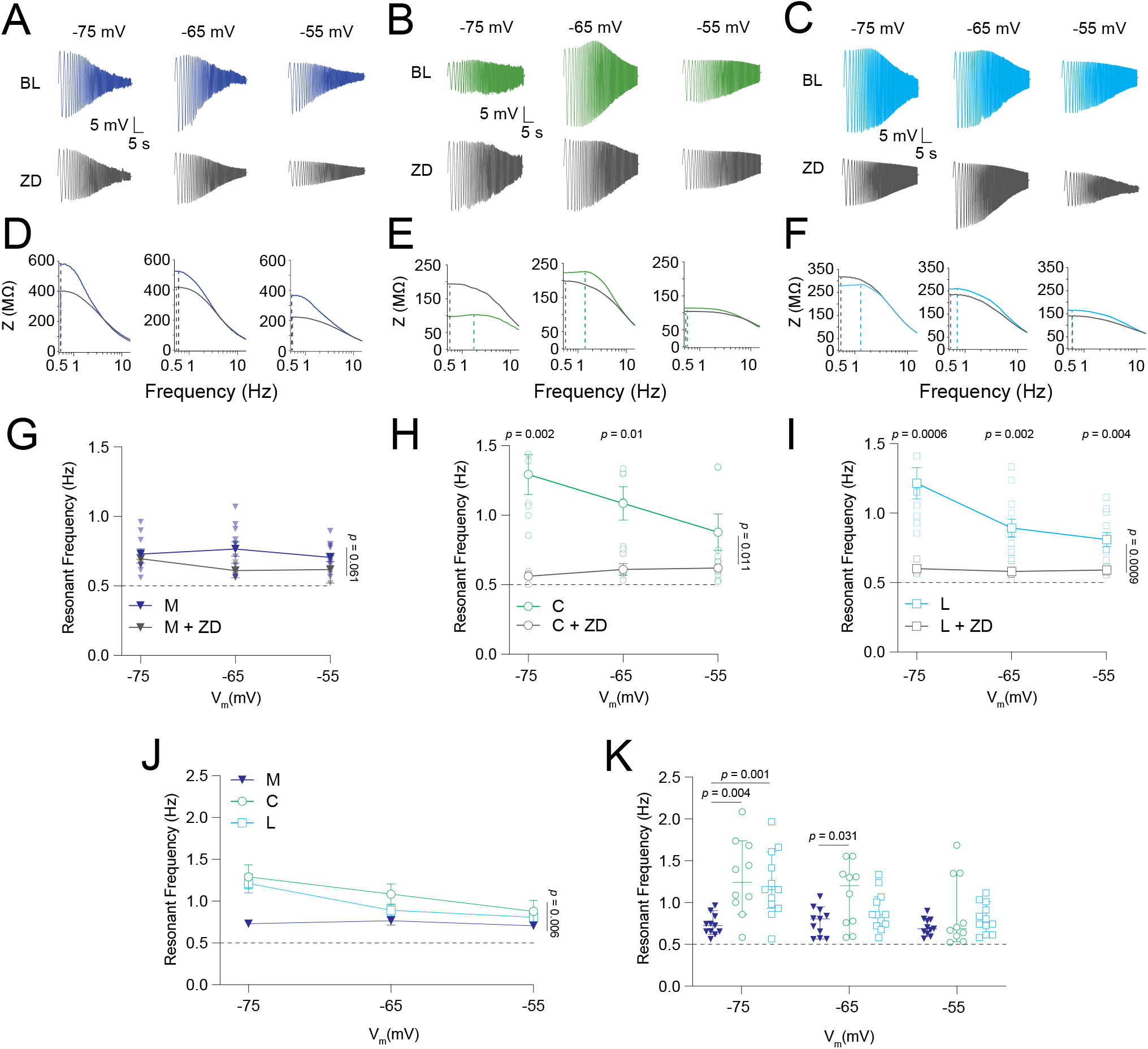
Subthreshold Chirp properties across MD subnuclei. **A**. Representative traces showing voltage responses from M neurons to a Chirp current input of 0.5 to 20 Hz while held at -75 mV, -65 mV, and -55 mV before and after wash-on of ZD7288. **B**. Same as A with C neurons. **C**. Same as A with L neurons. **D**. Impedance plots for corresponding M traces from A. **E**. Impedance plot for corresponding C traces from B. **F**. Impedance plots for corresponding L traces from C. **G**. R_f_ data from M neurons before and after wash-on of ZD7288. Two-Way ANOVA, Sidak’s Multiple Comparisons. **H**. Same as G with C neurons. **I**. Same as G with L neurons. **J**. Average R_f_ in MD neurons across holding voltages. **For all data**. N(neurons/animals): M=11/7, C=10/6, L=12/9

#### MD neurons primarily express HCN subunits 2 and 4

Previous studies on the prevalence of HCN subunits within the MD have primarily used immunohistochemistry to identify HCN subunit expression (Notomi and Shigemoto 2004). This method provides insight into expressed protein within a brain region, however, the localization of HCN channels in dendritic compartments in some brain regions (Magee 1998; Kalmbach et al. 2015) makes it difficult to segregate HCN expression into specific thalamic subregions. Here we used *in situ* hybridization on coronal slices including MD using probes for HCN 1-4. As expected, subjective assessment of hippocampal HCN mRNA revealed a large amount of fluorescence associated with HCN 1 and 2, a moderate amount associated with HCN 3, and little to no labeling in association with the HCN 4 probe. We assessed the presence of mRNA in MD at bregma -1.58 mm. **Fig. 7A** shows the location of MD subnuclei as well as the results of probes for HCN 1-4 and GBX2, a protein critical for MD development that is indicated as a marker for L and M MD subnuclei (Onishi et al. 2022). Horizontal line scans measuring the fluorescence across MD were taken for each slice matching the indicated bregma levels and normalized to compare the relative intensity of each HCN subunit probe within MD. We found that of the 4 HCN subunits, HCN 2 and 4 were the most prevalent within MD (**Fig. 7B**). We used the hippocampus as a positive control for the HCN probes because HCN 1-4 are expressed in this region (Moosmang et al. 1999; Santoro et al. 2000; Dougherty et al. 2013) and it is present in all analyzed coronal slices including MD (**Supp. Fig. 2A**).

**Figure 7:**
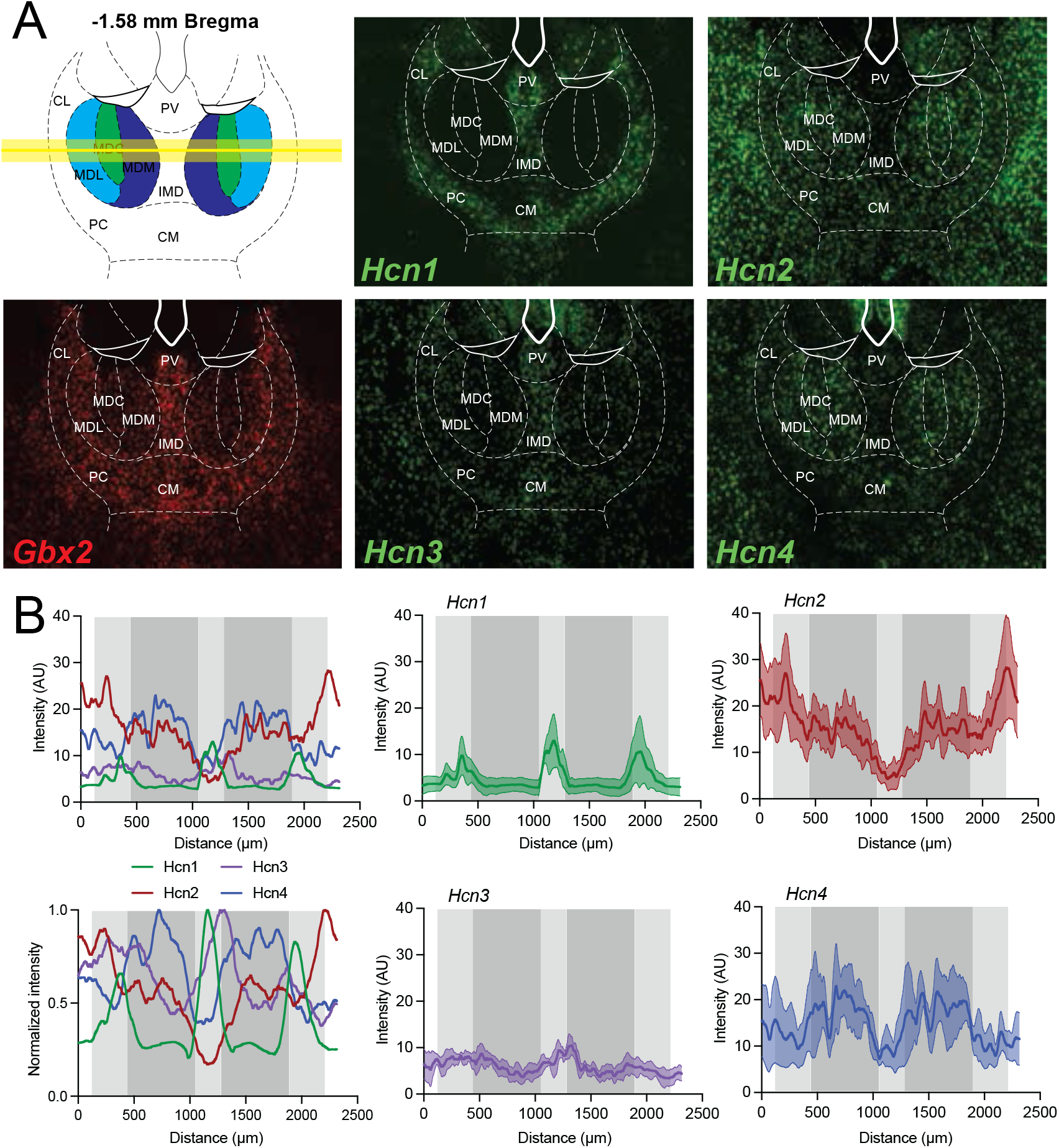
*In situ hybridization* reveals HCN2 and 4 as the primary HCN subunit mRNA present in MD neurons. *Hcn* mRNA prevalence across the mediodorsal thalamus (MD)—FISH experiments revealed the expression patterns of transcripts for HCN1, HCN2, HCN3, and HCN4. **A**. 20x Thunder images from MD at approximately bregma -1.58 with an anatomical map highlighting the MD subregions (top left) and *Gbx2* (bottom left) as a control marker for the region, *Hcn1* (top middle) is mostly visible in the CL and IMD with little to no signal in the MD. *Hcn2* (top right) is expressed in both the MD and outer portions of the thalamus and *Hcn3* (bottom middle) is expressed at low levels throughout with some signal concentrated in the IMD. While *Hcn4* (bottom right) is also found throughout the thalamus, it appears to show concentrated expression within the MD and lower expression in the CL and IMD. **B**. Quantification of signal across posterior sections of the thalamus (n=3) showing both individual channels and the combined data. Signal was smoothed using a Savitsky-Golay filter before being averaged. Individual plots of signal intensity show the mean intensity +/-SEM.

## Discussion

In this study we utilized multiple techniques to demonstrate how differences in HCN expression levels and activity uniquely affect MD subnuclei. We observed distinct levels of I_h_ in MD neurons from M, C, and L subnuclei using voltage clamp electrophysiology. Current clamp experiments directly preceding I_h_ measurement revealed intrinsic properties affected by the differences in HCN activity across subnuclei, primarily that greater HCN activity results in decreased R_N_, increased voltage sag, and an increase in current. Previous research specifically targeting thalamocortical neurons in the M and L region of MD identified differences in R_N_, tau, and action potential firing; properties associated with HCN activity (Lyuboslavsky et al. 2024). When compared across a range of voltages covering the physiological spectrum of thalamic neurons from alert states (∼ -55 mV) to restful states (∼ -75 mV), we found that current clamp properties associated with HCN activity, such as R_N_, membrane tau, and sag, varied based on resting membrane potential. Thalamic neurons exhibit state dependent firing properties depending on resting membrane potential (Llinás and Steriade 2006). It is critically important to understand how intrinsic neuronal properties, dictated primarily by the complement of voltage gated ion channels present in a neuron, interact membrane voltage. Differential levels of I_h_ across MD subnuclei may play a role in state switching or information processing. For instance, an M neuron that has relatively small I_h_, large R_N_, and a slower membrane tau will integrate incoming information over a broad time range, making it fire readily in response to incoming synaptic information across all physiological membrane potentials. Conversely, a C cell with relatively high I_h_, low R_N_, and a short membrane tau will require much greater synaptic activation and temporal coordination to reach action potential threshold when the membrane is hyperpolarized in the restful state (greater I_h_ activity) as compared to the same cell when the membrane has become depolarized to the alert state (reduced I_h_ activity).

Our data show that differential HCN expression in MD thalamus results in different R_N_ across M, C, and L MD subnuclei. R_N_ directly relates to input requirements to elicit an action potential or burst of action potentials from a neuron. Differing HCN expression in MD may be acting as a selectivity filter for information passing through neurons in each subnucleus. M neurons, which have relatively high R_N_, pass information easily whereas C neurons, which have relatively low R_N_, require a high degree of temporal synchrony for an action potential to be generated. Another aspect to consider in HCN control over neuronal sensitivity is the effect of cyclic nucleotides on HCN function. HCN 4 has greater sensitivity to the binding of cyclic nucleotides compared with HCN 2 (Wang et al. 2001; Wainger et al. 2001), indicating that signaling pathways involving cyclic nucleotide generation or suppression could vastly alter the intrinsic properties of MD neurons.

Using whole cell voltage clamp, we identified differing I_h_ among all subnuclei of MD. Additionally, we identified a negative shift in V_1/2_ in M neurons compared with C and L neurons. Although whole cell voltage clamp comes with inherent space clamp errors in the measure of kinetic events, studies in thalamic neurons using dual recordings in the soma and dendrites identify the neurons as being electrotonically compact with little signal loss identified between electrodes, particularly in soma to dendrite voltage transfer (Connelly et al. 2015). This data suggests a difference in HCN activity between M neurons and other subnuclei of MD. The two most parsimonious explanations would be differential levels of cyclic nucleotide activity and differing HCN subunit expression levels. To identify the HCN subunits expressed in MD, we used *in situ* hybridization to test for HCN 1-4. Previous studies have used immunohistochemistry to investigate HCN subunits present in MD (Notomi and Shigemoto 2004). We opted for a soma-restricted technique for more reliable localization of fluorescent signal to the region of interest. Our results suggest that HCN 2 and 4 are the most prevalent subunits across MD neurons. We used GBX2 to confirm MD identity since both M and L cells have GBX2 mRNA (Onishi et al. 2022). However, this signal was not distinct across the subnuclei of MD to allow for the colocalization and separation of subnucleus HCN expression.

Others have looked at molecularly and synaptically identified populations of neurons in MD (Onishi et al. 2022; de Kloet et al. 2021; Lyuboslavsky et al. 2024; Ordemann et al., n.d.). Here, we observed physiologically and anatomically defined subnuclei. Thus, we cannot speculate on the overlap of the cell populations we recorded from and those reported previously. Additionally, these experiments were performed based on the subnucleus location of the neuron, meaning that the specific connectivity of neurons recorded in this study is unknown. We provide here a physiological method of MD neuron identification that, in conjunction with anatomical localization, supports the accurate identification of MD subnucleus to which a neuron belongs. Further investigations will focus on identifying how differences in I_h_ contribute to the processing of information and the functional output of MD neurons.

Rhythmic activity is a hallmark of thalamic nuclei (Llinás and Steriade 2006). Oscillatory activity within the thalamus has been shown to align rhythmic activity throughout much of the brain (Lakatos et al. 2019; Schreiner et al. 2022) and is associated with complex cognitive tasks such as working memory (Fuster and Alexander 1971; Alexander and Fuster 1973; Bolkan et al. 2017; Parnaudeau et al. 2013) and social behavior (Ferguson and Gao 2018; Ramesh et al. 2025). HCN channels have been identified in the production of rhythmic activity in thalamic neurons (Llinás and Steriade 2006; Zobeiri et al. 2018; 2019; Kanyshkova et al. 2012). The presence of HCN in a neuron amplifies incoming signals at a specific R_f_ and, when expressed dendritically, temporally aligns synaptic signals in the theta to gamma range at the soma (Narayanan and Johnston 2008; Vaidya and Johnston 2013). We found that MD subnuclei with greater I_h_ also showed an increased R_f_. Interestingly, C and L MD neurons displayed the greatest resonant frequencies; C and L neurons also showed the most depolarized I_h_ V_1/2_ values. Since HCN4 has the most hyperpolarized V_1/2_ of the HCN subunits, this could indicate differential subunit expression across MD subnuclei, with M cells expressing greater HCN4 and C and L neurons expressing greater HCN2, accounting for more depolarized V_1/2_ values. The slower time course of activation and hyperpolarized V_1/2_ of HCN4 subunits (Wang et al. 2001; Wainger et al. 2001) could make R_f_ measurements difficult to obtain based on slower kinetics and reduced voltage sensitivity. Understanding this limitation of the linear chirp stimulus, we used a logarithmic chirp stimulus to bias the stimulus to gather more cycles at lower frequencies. We found that both C and L cells showed R_f_ values that deviated from controls using ZD7288 to block I_h_, indicating that M and L MD neurons show resonance at the measured membrane voltages. However, M neurons revealed no deviation between ZD7288 treated and baseline recordings indicating a lack of resonance. The distinction between resonant (C and L) and non-resonant (M) MD neurons further solidifies the view that M neuron intrinsic properties are stable across membrane voltage. This suggests that even at hyperpolarized membrane potentials, M neurons may show greater sensitivity to incoming information while C and L MD neurons likely require greater signal specificity, or temporal coordination, to elicit suprathreshold activity.

Neuronal R_f_ is heavily influenced by HCN activity; however, many voltage-gated ion channels widely expressed in neurons also interact to modulate R_f_ (Mishra and Narayanan 2025). The kinetics of Ca_v_3 activation and inactivation can impart resonance in neurons, particularly when expressed in conjunction with HCN channels (Suzuki and Rogawski 1989; Llinás and Steriade 2006). In MD neurons, Ca_v_3 channel expression causes the generation of low threshold calcium spikes, making the measurement of subthreshold oscillatory activity difficult without the blockade of voltage-gated calcium channel activity. In this study we focused on R_f_ in MD neurons imparted primarily by HCN channels through the comparison of R_f_ values before and after the use of ZD7288. Understanding how other voltage-gated ion channels contribute to resonance in highly rhythmic thalamic neurons will be critical to understanding thalamic rhythm generation and disorders involving thalamic dysrhythmia.

In many neurodevelopmental and neurological disorders, changes in HCN have been identified as a key driver in neuronal disease phenotype (Kim et al. 2018; Brager et al. 2012; Arnold et al. 2019; Chan et al. 2011). Our findings show subnucleus-specific differences in HCN expression in MD. This data, coupled with an established link between MD dysfunction and neurological disorders such as epilepsy (Kark et al. 2021), schizophrenia (Anticevic and Halassa 2023), and ASD (Brumback et al. 2018; Ordemann et al., n.d.), support the potential for HCN channels as a target for the treatment of neurological and neurodevelopmental disorders. Recent findings show a subnucleus specific decrease in HCN activity in the L region of MD in the neurodevelopmental disorder Fragile X syndrome (Ordemann et al., 2025). Decreased HCN could affect the ability of a specific subnucleus to filter out signals that would not be salient under normal conditions. Drugs targeted toward HCN channels are currently in use and have been investigated for their effects on neuronal activity. Lamotrigine is used to treat epilepsy by decreasing neuronal activity through the upregulation of HCN channels (Poolos et al., 2002) and was shown to treat NF1, a neurogenetic disorder, by restoring normal levels of HCN1 activity in a mouse model (Omrani et al., 2015). Our identification of the wide range of HCN activity across MD subnuclei provides a theoretical basis for the manipulation of HCN to treat neurological disorders and improve cognitive functions.

## Acknowledgments

This work was supported by NIMH R01 MH131857, the PERF Elterman research grant, the Phillip R. Dodge Young Investigator Award, the STARS award from The University of Texas System, startup funds from Dell Medical School, laboratory space from the College of Natural Sciences at UT Austin, The University of Texas System STARS award, and the Mulva Clinic for the Neurosciences at Dell Medical School at the University of Texas at Austin.

We thank Mendee Geist, Arwen Harris, David Nguyen, and Nyx Halder for their technical assistance. We thank members of the Brumback and Howard labs (especially MacKenzie Howard) for helpful discussions.

---

**Supplementary Figure 1:**
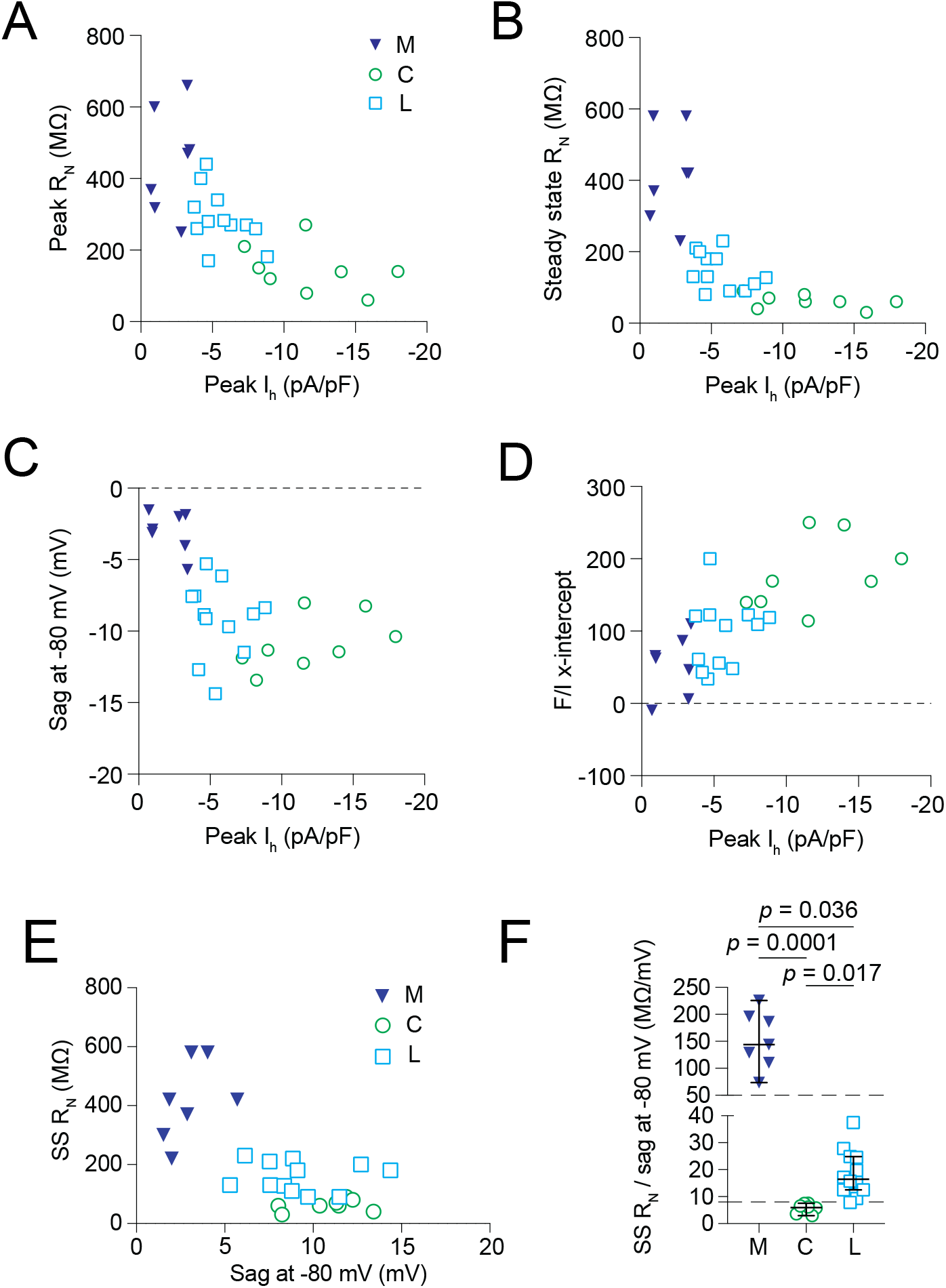
Current clamp properties used in K-means clustering plotted against I_h_. **A**. Peak R_N_ plotted against peak I_h_. **B**. Steady-state R_N_ plotted against peak I_h_. **C**. Voltage sag at -80 mV plotted against peak I_h_. **D**. X-intercept of frequency/current curves for each subnucleus plotted against peak I_h_. **E**. Scatter plot showing the relationship of SS RN to sag measured at a peak hyperpolarization of -80 mV across all subnuclei. **F**. Group data for M, C, and L neurons when the ratio between SS RN and sag at -80 mV was calculated. Kruskal-Wallis.

**Supplementary Figure 2:**
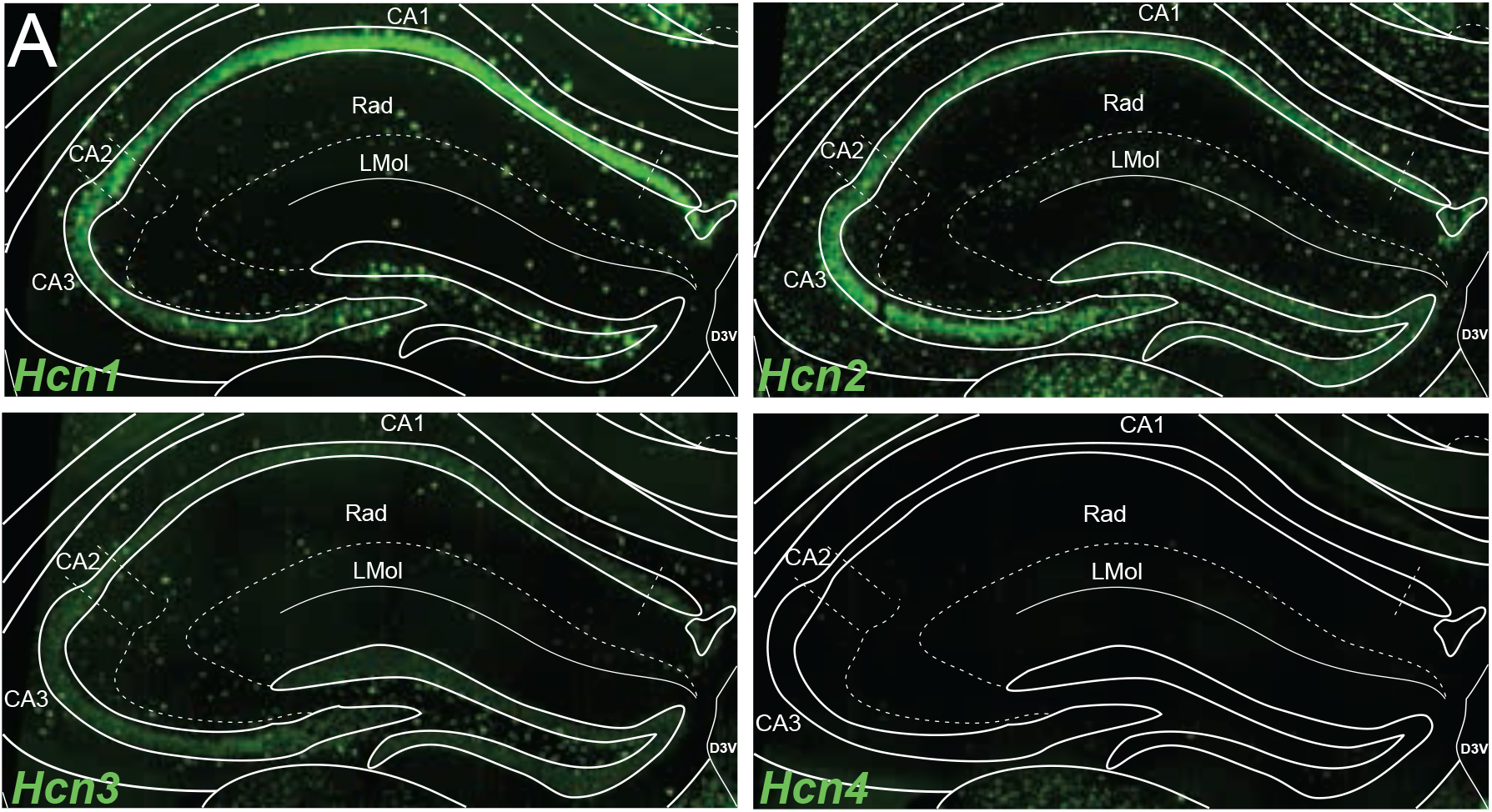
Positive control for *in situ* hybridization Hcn probes. **A**. In the hippocampus, *Hcn1* signal is strong within the pyramidal cell layer (CA1-CA3) and visible in interneurons but not as highly expressed in the dentate gyrus (DG). *Hcn2* is more evenly distributed while *Hcn3* is sparse and *Hcn4* does not appear to be expressed in the hippocampus consistent with existing data.

## KEY RESOURCES TABLE

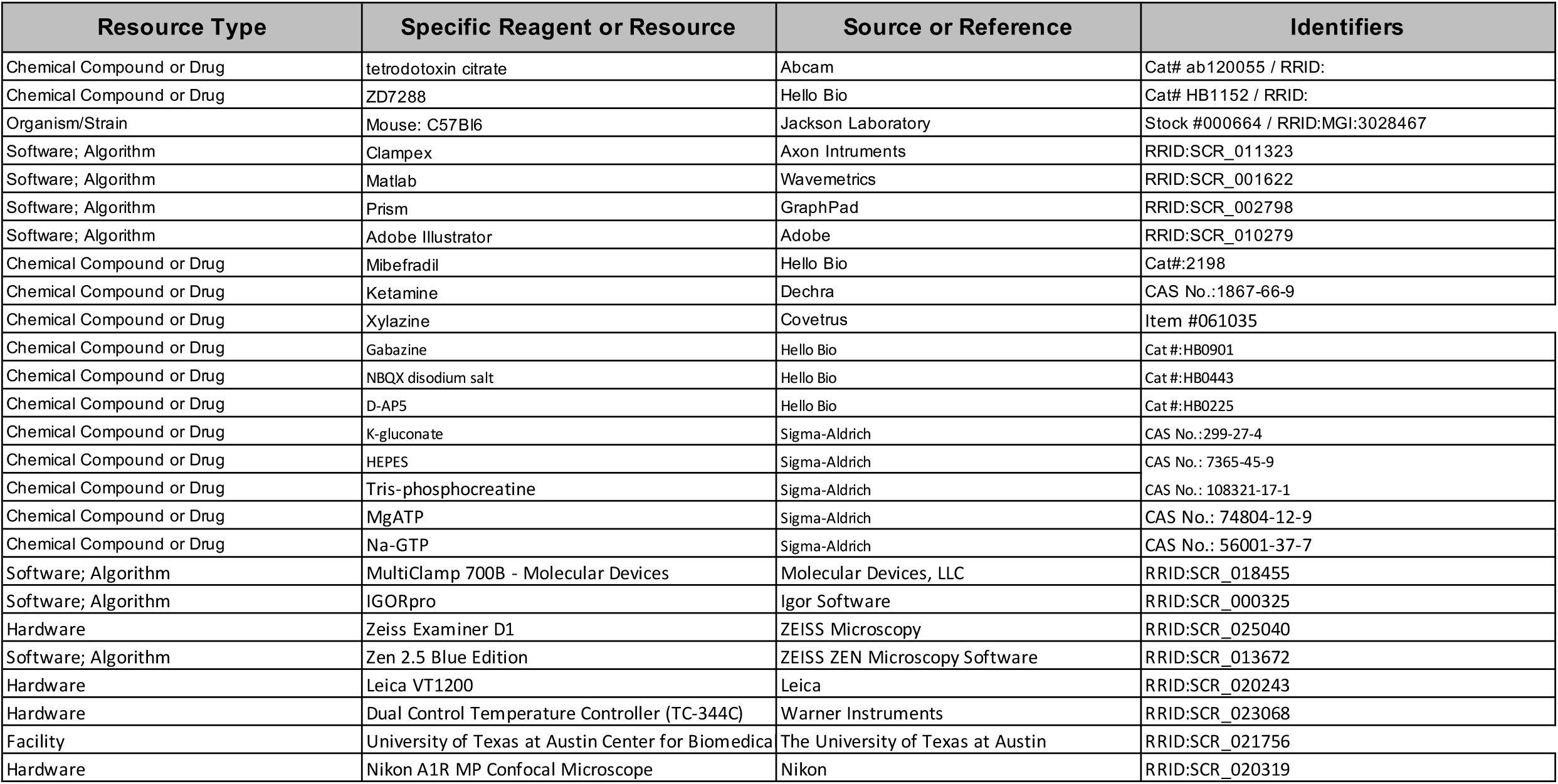

## Notes

Conflict of interest: The authors declare no competing financial interests.

### Competing Interest Statement

The authors have declared no competing interest.

